# SspA is a transcriptional regulator of CRISPR adaptation in *E. coli*

**DOI:** 10.1101/2024.05.24.595836

**Authors:** Santiago C. Lopez, Yumie Lee, Karen Zhang, Seth L. Shipman

**Affiliations:** Gladstone Institute of Data Science and Biotechnology, San Francisco, CA, USA; Graduate Program in Bioengineering, University of California, San Francisco and Berkeley, CA, USA; Department of Bioengineering and Therapeutic Sciences, University of California, San Francisco, CA, USA; Chan Zuckerberg Biohub – San Francisco, San Francisco, CA

## Abstract

The CRISPR integrases Cas1-Cas2 create immunological memories of viral infection by storing phage-derived DNA in CRISPR arrays, a process known as CRISPR adaptation. A number of host factors have been shown to influence adaptation, but the full pathway from infection to a fully integrated, phage-derived sequences in the array remains incomplete. Here, we deploy a new CRISPRi-based screen to identify putative host factors that participate in CRISPR adaptation in the *E. coli* Type I-E system. Our screen uncovers a novel host factor, SspA, which transcriptionally regulates CRISPR adaptation. One target of SspA is H-NS, a known repressor of CRISPR interference proteins, but we find that the role of SspA on adaptation is not H-NS-dependent. We propose a new model of CRISPR-Cas defense that includes independent cellular control of adaptation and interference by SspA.

## INTRODUCTION

CRISPR-Cas is an adaptive immune system found in archaea and bacteria, used to defend the host from foreign invaders, such as viruses or mobile genetic elements^1–5^. This defence is mediated by Cas (CRISPR associated) proteins, which are capable of creating immune memories of invading nucleic acids and using those memories to mount RNA-guided degradation of invaders in the event of a future encounter^6–9^. This process of storing immunological memory is known as CRISPR adaptation, and is mediated by a phylogenetically-conserved duo of proteins, Cas1 and Cas2, that form an integrase complex capable of inserting new DNA fragments (prespacers) into the cell’s CRISPR array^10–13^.

Studies spanning the past two decades have uncovered substantial mechanistic understanding of how CRISPR adaptation works and some of the key host factors that assist the CRISPR Cas1-Cas2 integrase complex in creating immune memories^14–24^. Double-stranded DNA fragments are the preferred substrate for the CRISPR Cas1-Cas2 integrases and can arise from a variety of sources, such as foreign DNA degradation by helicase-nuclease enzymatic complexes like the RecBCD complex^25^ or AddAB^26^ as well as from the replicating bacterial and phage genomes^27^. Fragments captured by the CRISPR Cas1-Cas2 integrases can then undergo trimming by Cas4^28^, DnaQ^29^ or other host exonucleases^30^, generating free 3’ OH groups required as substrates for spacer integration^21,29^. Cas1-Cas2 integrase docking at the Leader-Repeat junction of the CRISPR array requires the Integration Host Factor (IHF)^31–34^, which generates a bend in the Leader sequence that accommodates the integrase complex and allows it to form stabilising contacts with the DNA^33^. Docking enables the CRISPR integrase complex to catalyse a series of two nucleophilic attacks and add a new spacer at the Leader-Repeat junction^12,22,33^.

Spacer integration creates staggered double strand breaks at either end of the duplicated Repeat. Recent *in vitro* evidence suggests that host polymerases, in coordination with genome replication or transcription, could aid in repairing the CRISPR array^22^ (**Fig 1a**). The expanded and repaired CRISPR array is capable of supporting further rounds of spacer acquisition.

**Figure 1.**
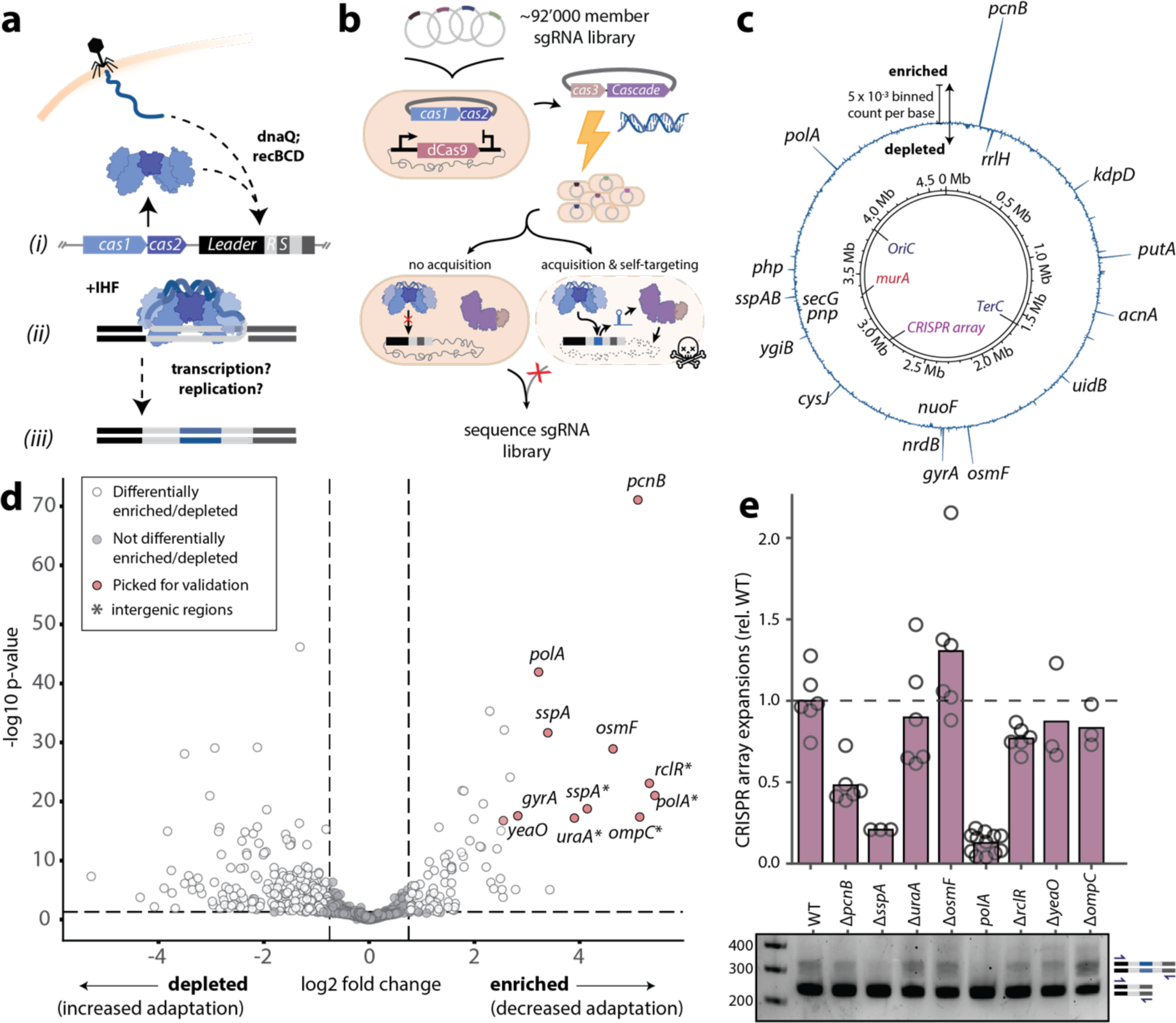
CRISPRi screen identifies adaptation host factors. **a.** Overview of the CRISPR adaptation process, highlighting key known host factors. **b**. Schematic of the CRISPRi adaptation host factor screen. **c**. Binned coverage plot of sgRNAs across the *E. coli* genome. sgRNA occupancy was calculated as the difference between the normalised (post/pre-screen) binned sgRNA counts per base of the experimental (+dCas9) and paired control (–dCas9) conditions. Regions of the genome with high (“enriched”) sgRNA coverage are interpreted to be genomic loci that positively regulate CRISPR adaptation; regions of the genome with low (or negative, i.e., “depleted”) sgRNA coverage are interpreted to be genomic loci that negatively regulate CRISPR adaptation. The highest-ranking regions with attributable genes are labelled; other labelled loci are the *Ori* and *Ter* regions, the *murA* gene, and the CRISPR-II array. *n* = 9 biological replicates. **d**. Volcano plot showing log2 fold change for each sgRNA versus adjusted –log10 p-values (*n* = 9 biological replicates). The horizontal dashed line represents an adjusted p-value of 0.05; the vertical lines represent log2 fold changes of –0.75 and 0.75. Genes targeted by sgRNAs differentially enriched that were selected for individual validation are coloured in pink. **e**. Top: deep-sequencing based measurement of the rates of new spacer acquisition in Keio knockouts harbouring pSCL565, after growth for 48h in liquid culture without induction of Cas1-Cas2 expression. Acquisition rates are shown relative to the wild-type parental strain. Open circles represent biological replicates (*n* ≥ 3), bars are the mean (one-way ANOVA effect of strain *P*<0.0001; Sidak’s corrected multiple comparisons for wild-type vs. knockouts, Δ*pcnB* P=0.00217, Δ*sspA* P=0.000102, *polA* ΔKlenow P<0.0001; others ns). Bottom: representative agarose gel for the data shown. Expansions of the CRISPR array can be seen as higher sized bands above the parental array length. Additional statistical details in **Supplemental Table 1**.

Despite this knowledge, several open questions remain, including what host factors are responsible for regulation of CRISPR-Cas activity^35–39^, and repair of the CRISPR array post-spacer integration^22,40^. Furthermore, in contrast to noteworthy successes in heterologous reconstitution and harnessing of the CRISPR interference machinery across the tree of life, most notably CRISPR-Cas9, there are conspicuously few reports of successful heterologous expression of a CRISPR adaptation system outside of its native host^41,42^, and no reports in eukaryotic systems. We, therefore, set out to discover additional host factors required for CRISPR adaptation.

Here, we develop and use a CRISPRi-based genetic screen to identify new host factors that participate in CRISPR adaptation in the Type I-E *E. coli* system. We report that a novel host factor, SspA, acts as a transcriptional-level regulator of CRISPR adaptation. We further find that SspA regulation of CRISPR adaptation does not function via H-NS, a known regulator of CRISPR interference and a member of the SspA regulon. Our data supports independent pathways for regulating the adaptation and interference components of CRISPR immunity, both downstream of SspA.

## RESULTS

### CRISPRi screen identifies adaptation host factors

We designed a genome-wide CRISPRi screen to identify potential host factors that participate in Type I-E CRISPR adaptation (**Fig 1a**). This screen utilizes a library of 92,919 gRNAs that are distributed across a population of *E. coli*, each of which direct a catalytically dead Cas9 (dCas9) to knock down transcription at a single locus, with multiple redundant gRNAs per gene^43,44^. We utilized this CRISPRi library in a negative selection scheme designed to deplete adaptation-competent cells. Specifically, we electroporated oligonucleotide prespacers that matched an essential gene into *E. coli* expressing a Type I-E CRISPR system. Integration of this prespacer into the CRISPR array would lead to the generation of a self-targeting crRNA and ultimately death of the adaptation-competent library members. CRISPRi knockdown of host factors involved in CRISPR adaptation would reduce adaptation and subsequent self-targeting, leading to enrichment of host factor gRNAs in the population following selection (**Fig 1b**).

For this screen, we used *E. coli* LC-E75, a derivative of K-12 MG1655, which encodes a Tetracycline-inducible dCas9 cassette integrated at the Phage 186 attB site^43^. We built an LC-E75 strain that carried a plasmid-encoded, IPTG-inducible Cas1-Cas2 cassette (plasmid hereon referred to as pSCL565). We then electroporated this strain and *E. coli* K-12 MG1655 (parental strain serving as a control) with a library of 92,919 plasmid-encoded sgRNAs, which target both coding and non-coding regions across the *E. coli* genome^43^. The libraries were grown overnight, and subsequently passaged and grown to mid-log phase (∼3h) with dCas9 induction; the remainder of the overnight library cultures were harvested for sgRNA library sequencing (pre-screen library). Then, cells were co-electroporated with (1) a plasmid encoding an m-Toluic acid-inducible *E. coli* Cas3-Cascade, the effector of the Type I-E CRISPR interference system, and (2) a 35bp dsDNA spacer targeting the essential gene *murA*. Cells were rescued in media containing inducers for Cas3-Cascade and dCas9 and antibiotics to select for their respective plasmids, cultured overnight, and harvested for sgRNA library sequencing (post-screen library). We extracted the sgRNA plasmid libraries and prepared samples for sequencing by amplifying the sgRNAs using a primer pool targeting the region upstream of the sgRNA promoter and downstream of the tracrRNA. The primers contained Illumina adapters to make the amplicons compatible with our downstream sequencing prep. Sequencing of the sgRNAs libraries yielded sgRNA counts for the dCas9-expressing LC-E75 and control dCas9-less parental strains, which allowed the calculation of the binned enrichment/depletion of sgRNAs across the *E. coli* genome (**Fig 1c**).

We found peaks of sgRNA enrichment that were distributed across the *E. coli* genome and did not cluster around the *murA* locus. Additionally, we identified *polA*, *priA* and *gyrA*, essential genes previously suggested to play a role in the CRISPR adaptation process. This highlights the advantage of a knock-down approach over transposon-based knock-out approaches, where essential genes would have been lost from the library altogether. We found several other regions of the *E. coli* genome where sgRNAs were strongly enriched, suggesting additional host factors.

We quantified differentially enriched or depleted sgRNAs from their cumulative sgRNA counts (sum of all sgRNAs per gene), by comparing each experimental sample (+dCas9) to its paired control (–dCas9) using PyDESeq2 package^43–45^. We filtered out genes with less than 10 cumulative reads, and controlled for variation in relative sgRNA library composition by including pre-screen sgRNA counts as an interaction factor in the model. We found 571 differentially enriched/depleted genes and gene-adjacent regions, out of a total of 12,809 gene/gene-adjacent regions considered in our analysis (**Fig 1d**). Interestingly, a subset of the differentially enriched genes (i.e., CRISPR adaptation deficient when knocked-down) also had their gene-adjacent regions differentially enriched (shown with asterisked gene names).

We selected the top 8 gene regions with highest log_2_ fold changes for individual validation using knockout mutants from the Keio collection^46^ in a naive adaptation assay. One additional gene, *gyrA*, is essential and could not be validated with a knockout. Although polA knockouts are non-viable, a *polA* Klenow fragment deletion mutant is viable^47^, and was thus used in validation assays alongside the other non-essential genes.

We electroporated wild-type and knockout strains with pSCL565 and grew them in liquid culture for 48h without inducers for Cas1-Cas2 to achieve a moderate level of expression from transcriptional leak. We then sequenced the CRISPR II array of these cells (i.e., endogenous CRISPR array flanked by the *ygcE* and *ygcF* genes^48^, hereon referred to as CRISPR-II) and quantified the rate of CRISPR adaptation as the fraction of sequenced arrays that had acquired new spacers. Biological replicates run on different days were normalised to the CRISPR adaptation rate of the wild-type parental Keio strain (**Fig 1e**). We found that 3 mutants showed significantly decreased rates of CRISPR adaptation compared to the wild-type strain: *pcnB*, *sspA*, and *polA Δ*Klenow.

### Features of spacers acquired in Knockout Strains

Spacers captured by the CRISPR Cas1-Cas2 integrases come from a variety of sources. Defence associated sources include mobile genetic elements and phages. However, in the absence of interference machinery, spacers derived from the bacterial genome and plasmids accumulate (**Fig 2a**). We next tested whether any of the hits that we chose for validation modified the source of new spacers. We found that, consistent with previous findings^14,49^, the majority of new spacers acquired in the wild-type strain were plasmid-derived (**Fig 2b**). This finding held for all mutants except *polA Δ*Klenow, which acquired spacers solely from the genome. The breakdown of spacer origin as a percent of all newly acquired spacers starkly illustrates this finding (**Fig 2c**).

**Figure 2.**
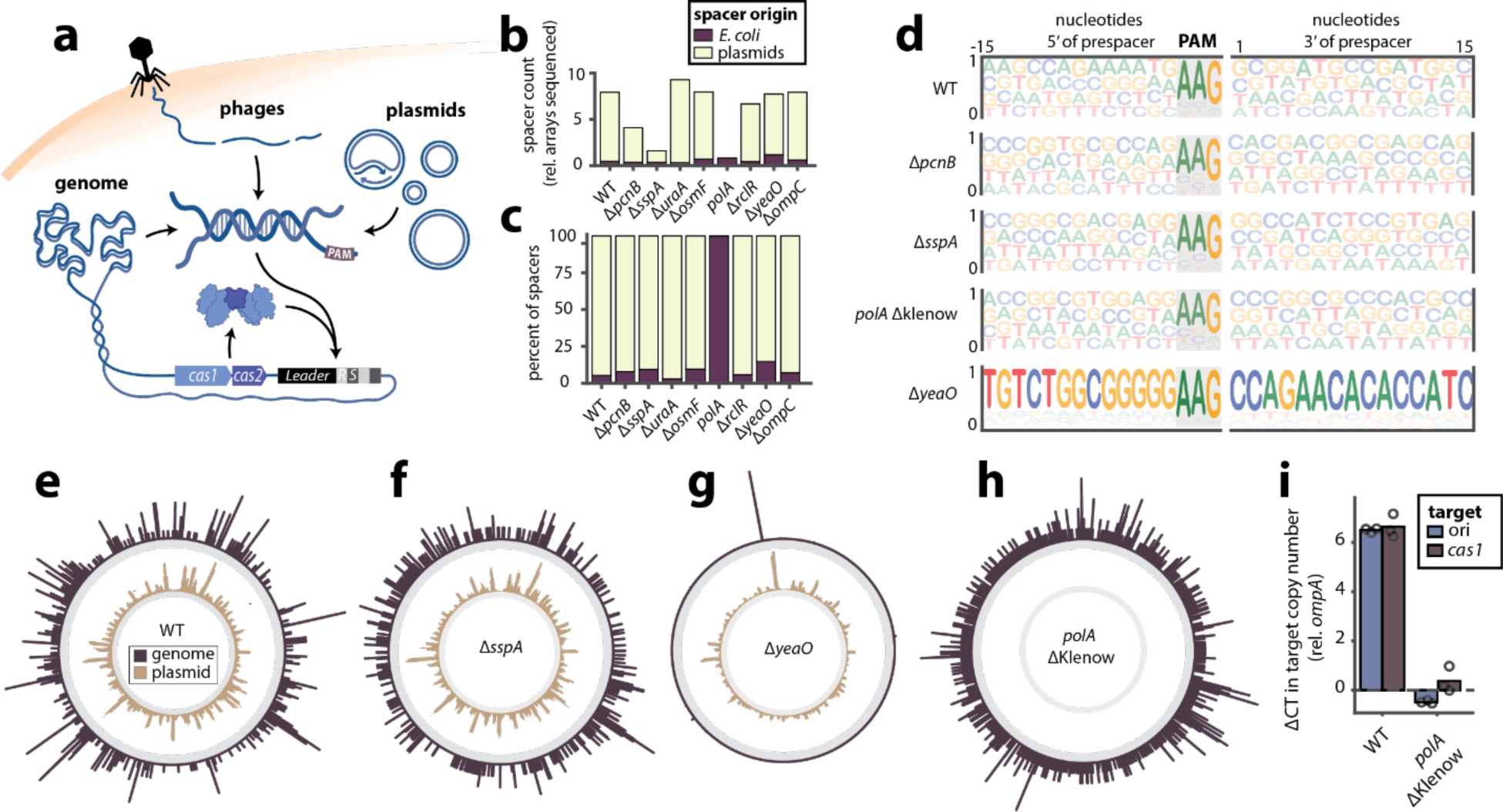
Features of spacers acquired in knockout strains. **a.** Prespacer substrates for CRISPR adaptation arise from a variety of sources. **b.** Breakdown of normalised spacer count (total number of new spacers / number of CRISPR arrays sequenced) according to spacer origin (*E. coli* or plasmid) and strain of interest. **c.** Breakdown of percent of spacer attributable to each spacer origin (*E. coli* or plasmid) and strain of interest. **d.** Motifs in the 15bp up- and downstream of the newly acquired spacer in its source location. **e-f:** Binned coverage plot of newly acquired spacer across the *E. coli* genome (outer, purple) and pSCL565 plasmid (inner, tan) for the wild-type strain (**e**) and derivatives (**f-h**). See **Extended Data figure 1** for the full set. **i.** qPCR-based measurement of the relative copy number of pSCL565 *Ori* and *cas1* sequences in the wild-type and *polA* ΔKlenow mutant. Open circles represent biological replicates (*n* ≥ 3), bars are the mean (one-way ANOVA effect of strain and target *P*<0.0001; Sidak’s corrected multiple comparisons for wild-type vs. Δ*sspA*, CDF ori copy number P<0.0001, *cas1* copy number P<0.0001). Additional statistical details in **Supplemental Table 1**.

We next sought to determine whether the differences in new spacer acquisition could be explained by a change in PAM preference or other motifs up- or downstream of the spacer. We searched 15bp up- and downstream of the newly acquired spacer in its source location, and found that all mutants showed similar PAM preferences to the wild-type strain, consistent with previous reports^50^. Similarly, all mutants except the *yeaO* deletion mutant showed no additional up- and downstream motif preferences of preference, beyond the AAG PAM.

The *yeaO* mutant displayed strong motif preferences up- and downstream of the genome-derived spacers (**Fig 2d**), which prompted us to map all newly acquired spacers for each mutant to their respective source on either the *E. coli* genome or pSCL565 plasmid (**Figs 2e-h**; see **Extended Data Figure 1** for an expanded view of these figures). We found that the distribution of new spacers from both sources were mostly consistent between the wild-type and the mutants tested (**Extended Data Figure 1a-i)** with two exceptions: the *yeaO* and *polA Δ*Klenow mutants.

We found that, as suggested by the prespacer neighbourhood motif analysis (**Fig 2d**), the *yeaO* mutant acquired spacers almost uniquely from one location in the genome, which maps to the gene *insG*, encoding an IS4 transposase (**Fig 2g**; **Extended Data Figure 1h**). It is possible that the high number of *insG*-derived spacers were a product of an early acquisition event from which most of the sequenced arrays descended, or if there had been multiple independent acquisition events leading to *insG*-derived spacers across CRISPR arrays. We can distinguish the two by looking at multiply expanded CRISPR arrays, or arrays that had acquired two or more new spacers, due to the CRISPR Cas1-Cas2 integrases’ preference for Leader-proximal spacer insertion: this feature makes the CRISPR arrays temporally ordered, with spacers acquired at a later stage being closer to the Leader sequence than spacers acquired earlier, or than vestigial spacers^41,49,51,52^. We found *insG*-derived spacers in multiple different positions with respect to the Leader and found *insG*-derived spacers in arrays with distinct additional new Leader-distal spacers across three biological replicates (**Extended Data Figure 2**), suggesting that *insG*-derived spacers represent more than one acquisition event in parallel lineages.

Though there is no literature describing *yeaO*, it is predicted to contain a DUF488-like domain, which is ubiquitously distributed across prokaryotes and some viruses, and has been found in genomic neighbourhoods linked to prophages and defence islands^53^. Structural searches using DALI^54^ and Foldseek^55^ suggested that YeaO is structurally similar to putative transcriptional regulators; in turn, *insG* is predicted to encode a transposase in a IS4 transposable element, previously been reported to be non-mobilised in *E. coli*^56^. We suggest that YeaO could be regulating the InsG-mediated mobilisation of IS4; the transposition events could cause the generation of DNA fragments that could serve as prespacers for the Cas1-Cas2 integrases, decoying these away from the electroporated *murA* prespacer and phenocopying a decrease in CRISPR adaptation, thus explaining the detection of YeaO in our screen.

The *polA Δ*Klenow mutant had a similar distribution of prespacers originating from the genome when compared to the wild-type, but we were unable to map any prespacers to the plasmid, suggesting that the plasmid was unable to serve as a source of prespacers in this mutant (**Fig 2h**; **Extended Data Figure 1f**). Given the loss of CRISPR adaptation from the pSCL565 plasmid but wild-type levels of CRISPR adaptation from the genome (**Fig 2b**), we hypothesised that the *polA* ΔKlenow mutant could be deficient in plasmid replication^57^. To test this, we measured the relative number of copies of the pSCL565 *Ori* and *cas1* sequences (the latter also found in the genome) in the wild-type and *polA* ΔKlenow mutant. We found a nearly 80-fold difference in the relative number of copies of pSCL565 that the *polA* ΔKlenow mutant contains compared to the wild-type strain, which would explain this strain’s decreased ability to acquire new spacers from the plasmid (significantly decreased number of plasmid copies per cell) and also why it was identified as a hit in the initial screen (**Fig 2i**).

### SspA is a transcriptional regulator of CRISPR adaptation

Our CRISPR adaptation assays and downstream analysis of acquired spacers revealed that Δ*sspA* was consistently and significantly defective in naïve CRISPR adaptation, despite no other noticeable differences in the features of its acquired spacers when compared to the wild-type parental strain (**Fig 2b, c, e, and f**). Additionally, we found that the decrease in CRISPR adaptation in the Δ*sspA* background was not due to decreased protein expression levels from pSCL565 (**Extended Data Figure 3**). We thus selected *sspA* for further mechanistic characterisation.

*E. coli* SspA was discovered four decades ago during a screen for proteins induced by the stringent response^58^. Over the years, its reported cellular functions have increased, and SspA has become particularly linked to global stress response^59,60^ through its action as an RNA polymerase (RNAP)-associated protein^61,62^. Crystal structures of *E. coli* RNAP-promoter open complex with SspA have revealed that SspA inhibits σ^70^ promoter escape through contacts with both RNAP and σ^70^ through a conserved PHP motif^59,62–64^. This promoter escape inhibition induces a rewiring of the cellular transcriptomic landscape towards expression of σ^S^ genes, with implications on stress tolerance, motility and virulence^59,62–64^. The *sspA* gene is encoded in a two-member operon, upstream of *sspB*. SspB acts as a specificity-enhancing factor for the ClpXP protease^65^. It helps maintain protein homeostasis by escorting SsrA-tagged peptides, resulting from stalled ribosomes, to the ClpXP protease and promoting their degradation (**Fig 3a**), thus simultaneously freeing ribosomes and replenishing the pool of amino-acids that can become a precious resource in conditions of starvation.

**Figure 3.**
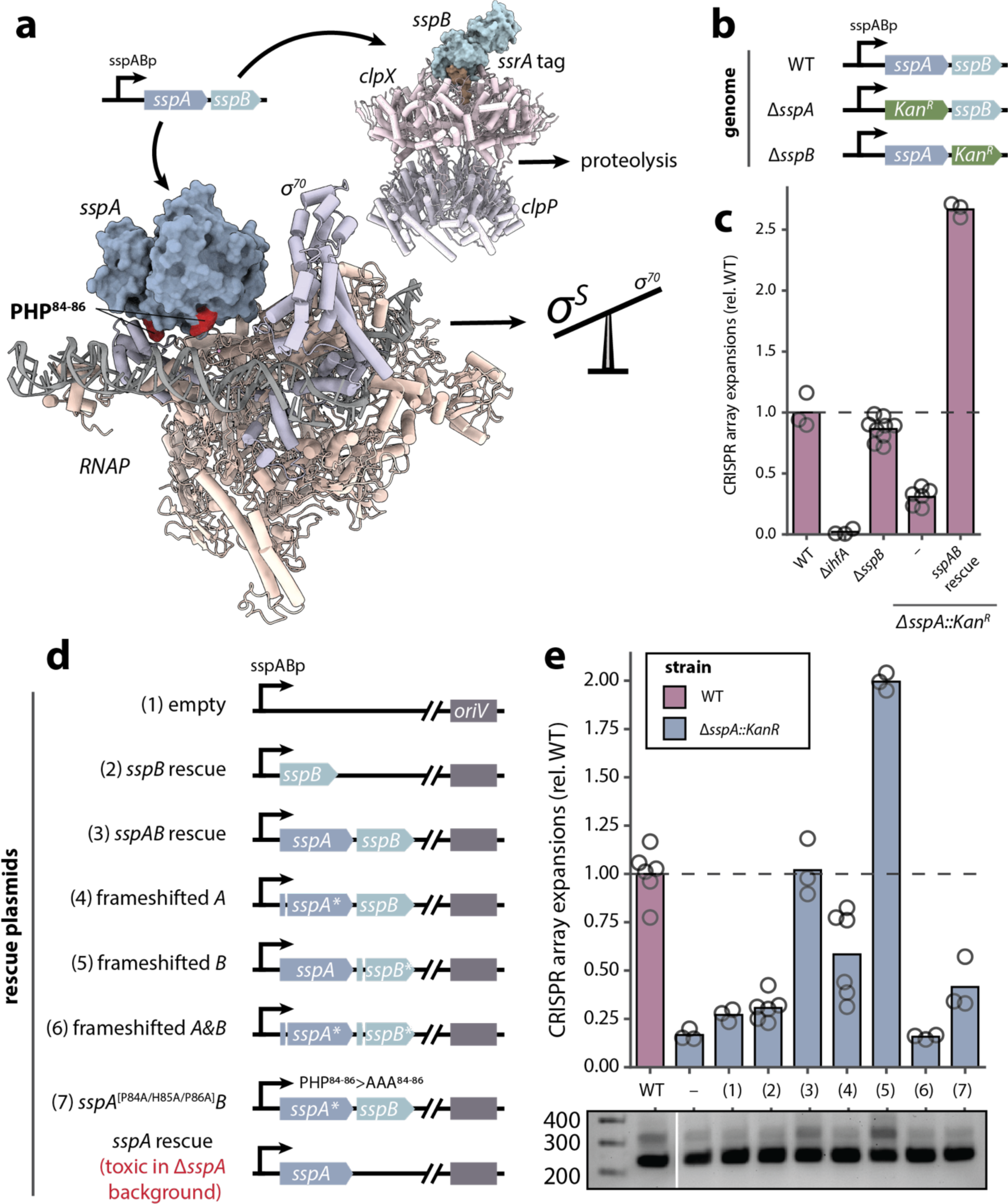
*sspA* is a transcriptional regulator of CRISPR adaptation. **a**. *sspAB* operon, proteins and function. Bottom left: crystal structure of an SspA dimer (blue) in complex with *E. coli* RNAP-promoter open complex, showing the conserved SspA PHP^84–86^ residues (red) interacting with RNAP (pink) and σ^70^ (purple) (PDB 7DY6^62^). Top right: crystal structure of SspB escorting an SsrA-tagged substrate being delivered to the ClpXP protease complex (PDB 8ET3^65^). **b**. Schematic of the *sspAB* operon of WT, Δ*sspA*::*kan^R^* and Δ*sspB*::*kan^R^* strains. *kan^R^*: kanamycin resistance cassette. **c**. Deep-sequencing based measurement of the rates of new spacer acquisition in strains harbouring pSCL565 and, in the case of the Δ*sspA*::*kan^R^*, either an empty plasmid or a low (∼5) copy plasmid encoding the *sspAB* operon, after growth for 48h in liquid culture. Adaptation rates are shown relative to the wild-type parental strain. Open circles represent biological replicates (*n* ≥ 3), bars are the mean. Horizontal dashed line represents the mean rate of spacer acquisition in the wild-type strain (one-way ANOVA effect of strain *P*<0.0001; Sidak’s corrected multiple comparisons for wild-type vs. knockouts, Δ*sspA* P<0.0001, Δ*sspB* P=0.109807; Δ*sspA* vs. Δ*sspB* P<0.0001). **d**. Schematic of the *sspAB* operon variant rescue plasmids. All plasmids are low (∼5) copy, and encode variants of the *sspAB* operon under its native regulation. Frameshift mutants of SspA (AN^5–6^>AQ^5–6^ GCC|AAC>GC**T**|*CAA*|C) and SspB (PR^9–10^>PS^9–10^ CCA|CGT>CCA|***T****CG*|T) encode sequences with single base insertions to cause protein translation to terminate early. The SspA PHP^84–86^>AAA^84–86^ mutant is RNAP-binding deficient and thus does not enable the shift in promoter use (α^70^ σ α^S^)^62^. A single *sspA* rescue plasmid yielded no transformants into the Δ*sspA*::*kan^R^* strain over multiple attempts. **e**. Top: deep-sequencing based measurement of the rates of new spacer acquisition in strains harbouring pSCL565 and, in the case of the Δ*sspA*::*kan^R^*, either an empty plasmid or a low (∼5) copy plasmid encoding variants of the *sspAB* operon as described in **d**., after growth for 48h in liquid culture. Adaptation rates are shown relative to the wild-type parental strain. Open circles represent biological replicates (*n* ≥ 3), bars are the mean. Horizontal dashed line represents the mean rate of spacer acquisition in the wild-type strain (one-way ANOVA effect of strain *P*<0.0001; Sidak’s corrected multiple comparisons for wild-type vs. knockouts, Δ*sspA* P<0.0001, Δ*sspA +* empty plasmid P<0.0001, Δ*sspA* + *sspAB rescue* P = 1, Δ*sspA + sspA*\* (PHP84-86>AAA84-86) & *sspB* rescue P<0.0001; Δ*sspA* vs. rescues, Δ*sspA* + empty vector P=0.997758, Δ*sspA* + *sspA*\* (PHP84-86>AAA84-86) & *sspB* P=0.334315, Δ*sspA* + *sspAB* P<0.0001, Δ*sspA* + *sspB* P=0.892991, Δ*sspA* + *sspA*\* & *sspB*\* (frameshifted) P=1). Bottom: representative agarose gel for the data shown. Expansions of the CRISPR array can be seen as higher sized bands above the parental array length. Additional statistical details in **Supplemental Table 1.**

Though we found *sspA* and not *sspB* as a significant hit in our screen, we sought to confirm that the defects in CRISPR adaptation observed in the Δ*sspA* mutant were due strictly to the lack of SspA, and not due to polar effects of this mutation on the downstream *sspB* gene. We compared the rates of CRISPR adaptation in wild-type strains to those in Δ*sspA::kan^R^* and Δ*sspB::kan^R^* mutants carrying pSCL565 (**Fig 3b**). We found that the Δ*sspA* mutant was deficient at new spacer acquisition, but the Δ*sspB* mutant acquired spacers at rates indistinguishable from the wild-type strain (**Fig 3c**). We attempted to deliver an *sspA* rescue plasmid into the Δ*sspA*::*kan^R^* strain, but this yielded no transformants over multiple attempts. However, we found that we could rescue the CRISPR adaptation phenotype to wild-type levels when Δ*sspA::kan^R^* carrying pSCL565 were additionally electroporated with an *sspAB* cassette, encoding the SspA and SspB proteins under control of their native promoter and on a low-copy (∼5) plasmid. This suggests that lack of *sspA* alone is sufficient to cause the loss of adaptation phenotype, and that this can be rescued by supplying a copy of the *sspA* gene in *trans*, under its native regulation.

Given these findings, we next sought to determine which part of the SspA protein was responsible for the loss of adaptation phenotype. We were particularly interested in the SspA PHP^84–86^ motif, which has been reported to be indispensable for stabilisation of interactions between SspA, α^70^ and the RNAP complex. Via this interaction, SspA acts as a transcriptional repressor of α^70^ promoters by inhibiting promoter escape^62^. Triple-Alanine substitutions in this motif cause pleiotropic cellular effects such as increased swarming and defects in acid-resistance and phage P1 growth^60,66^. Thus, given SspA’s role as a transcriptional rewiring agent, we decided to test whether an SspA PHP^84–86^>AAA^84–86^ mutant, deficient in α^70^-RNAP binding, would also phenocopy the Δ*sspA* mutant in terms of loss of CRISPR adaptation. To do this, we designed rescue plasmids, encoding variants of the *sspAB* operon under endogenous regulation, on low-copy (∼5) plasmids (**Fig 3d**).

We found that rescue plasmids encoding the full *sspAB* operon or an early frameshifted *sspB* could rescue CRISPR adaptation to levels comparable to wild-type. However, rescue plasmids encoding SspB, an early frameshifted *sspA*, early frameshifted *sspA* and *sspB*, and crucially, the SspA PHP^84–86^>AAA^84–86^-SspB RNAP binding mutant were all deficient in CRISPR adaptation (**Fig 3e**). Taken together, our data is consistent with a model in which SspA’s role as RNAP-σ^70^ interactor and transcriptional rewiring agent is required for functional CRISPR adaptation.

### H-NS regulates CRISPR interference downstream of SspA

Having established SspA’s role as a regulator of CRISPR adaptation, we sought to test whether it could also play a role in regulating CRISPR interference. As SspA has been shown to be rapidly and highly upregulated following lambda infection^67^, it could serve as a link between phage infection and CRISPR defence generally. Given previous reports of the role of SspA in downregulating levels of H-NS^60,68^, a repressor of the CRISPR interference machinery, we hypothesised that SspA could be acting on the CRISPR-Cas system via H-NS^69–71^ (**Fig 4a**).

**Figure 4.**
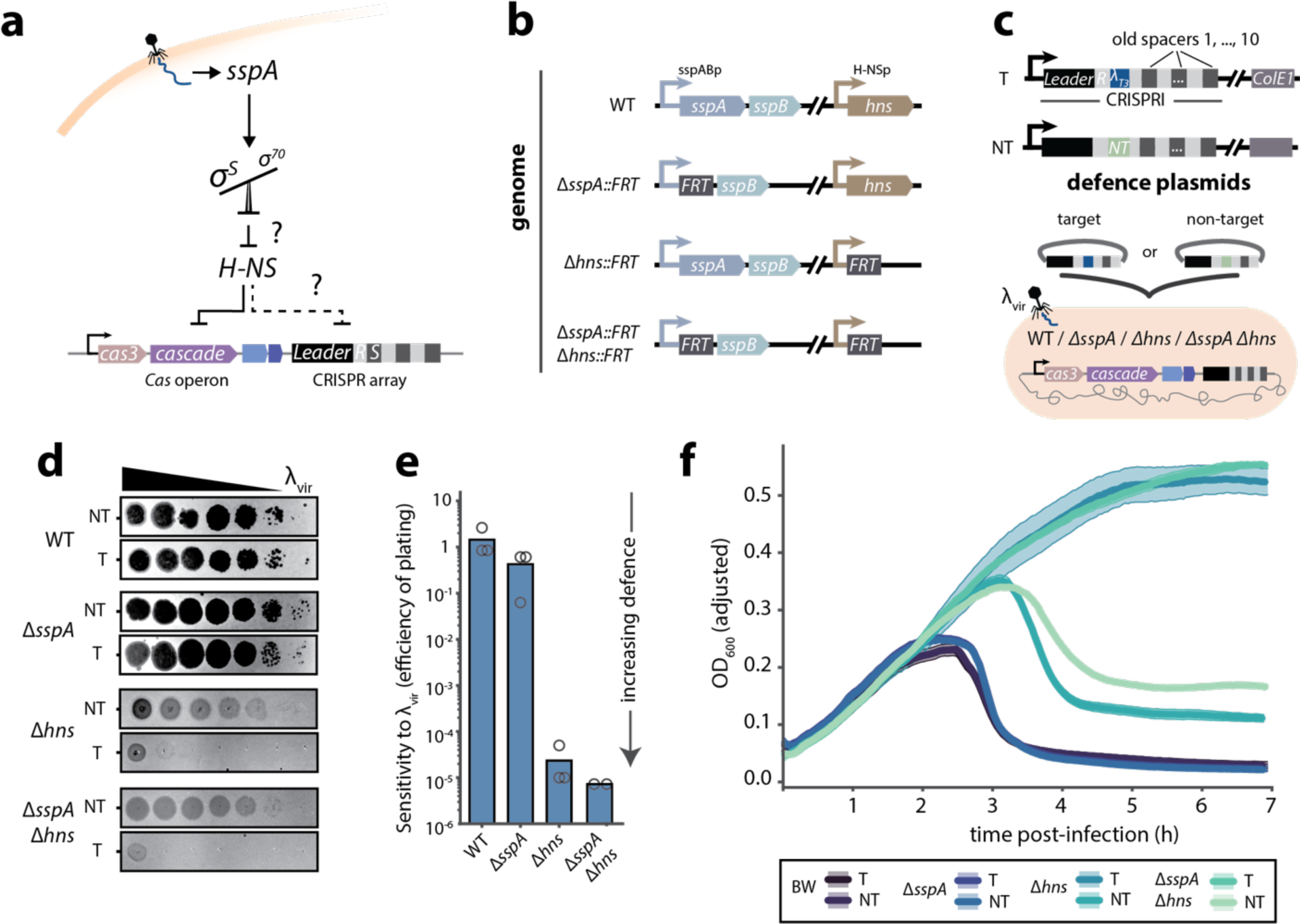
H-NS regulates CRISPR interference downstream of SspA. **a**. Model for SspA-mediated regulation of CRISPR-Cas defence. Phage infection triggers upregulation of SspA^67^, which in turn induces a global transcriptional shift towards 0*^S^*-regulated promoters. This results in H-NS downregulation^60,68^, induction of CRISPR-Cas mediated defence through de-repression Cas gene expression^69,70^, leading to increased rates of CRISPR adaptation and interference. **b**. Schematic of the *sspAB* and *hns* operons of WT, Δ*sspA*::*FRT,* Δ*hns*::*FRT* and Δ*sspA*::*FRT* Δ*hns*::*FRT* strains. *FRT*: flippase recognition target, a scar left after the removal of resistance cassettes. **c**. Schematic of the CRISPR interference-mediated defence assays in pre-immunised *E. coli* strains. Top: schematic of the CRISPR-I immunisation (defence) plasmids. All plasmids are low (∼5) copy, and encode an *E. coli* CRISPR-I array with a first spacer encoding either a Target (complementary to the α genome^1,70^), or a Non-Target (NT) spacer. Bottom: The experimental strains were electroporated with either the T or NT plasmid, and infected to varying titres of αvir. Note that the strains encode a complete endogenous *E. coli* Type I-E CRISPR-Cas system. **d**. Representative plaque assays of αvir on experimental strains (described above) pre-immunised with either T or NT defence plasmids. Strains were infected with αvir and grown on plates at 30°C for 16h. Full plaque assay plates for *n* = 3 biological replicates in **Extended Data Figure 4**. **e**. Efficiency of plating of αvir on experimental strains. Open circles represent biological replicates (*n* ≥ 3) of individual plaque assays, bars are the mean (one-way ANOVA effect of strain *P*=0.033454; Sidak’s corrected multiple comparisons for wild-type vs. knockouts, Δ*sspA* P=0.181757, Δ*hns* P=0.043319, *ΔsspA Δhns P* = 0.043316; for Δ*hns* vs. Δ*sspA Δhns* P=1). **f.** Anti-phage defence and growth in overnight liquid culture of experimental strains, post αvir infection (MOI: 0.1). Hue around solid line (mean) represents the standard deviation across 3 biological replicates.

To assess the effects of SspA on CRISPR mediated anti-phage defence and the potential interactions between SspA and H-NS in regulating this defence, we constructed Δ*sspA::FRT*, Δ*hns::FRT* and Δ*sspA::FRT* Δ*hns::FRT E. coli* strains (**Fig 4b**). Previous studies have shown that H-NS is a strong repressor of CRISPR-Cas gene expression, but that this repression can be relieved by knocking out H-NS^69,70^. This de-repression can result in defence against bacteriophages, provided that these cells’ CRISPR arrays encode one or more spacers targeting the phage genome (hereinafter referred to as “pre-immunised *E. coli*”)^1,69,70^.

We electroporated our mutant strains with a plasmid carrying a CRISPR array encoding a first spacer complementary to the lambda genome (T: target^1,70^) or a control CRISPR array with a non-target first spacer (NT: non-target). Then, we infected these pre-immunised strains with varying titres of α_vir_ and quantified phage defence (**Fig 4c**). Because of the pre-immunisation, this assay measures the ability of mutants to mount anti-phage defence via CRISPR interference and should be CRISPR adaptation-independent.

Plaque assays revealed that a wild-type strain was unable to mount defence against new rounds of infection even when pre-immunised with an anti-α spacer, as reported previously^1,69,70^ (**Fig 4d**). Pre-immunised Δ*sspA* mutants were similarly unable to defend against α_vir_. However, pre-immunised Δ*hns* mutants were capable of mounting considerable defence against α_vir_. Interestingly, we saw no differences in anti-α_vir_ defence between the Δ*hns* and Δ*hns* Δ*sspA* pre-immunised mutants, suggesting that the CRISPR-Cas mediated anti-phage defence observed in the Δ*hns* Δ*sspA* mutants was determined solely by the lack of CRISPR interference repression by H-NS, and that Δ*sspA* has no additive effect on CRISPR interference-mediated anti-phage defence on the Δ*hns* background. Quantification of efficiency of plating confirmed these findings (**Fig 4e**), as did additional experiments measuring anti-phage defence in overnight liquid culture growth assays (**Fig 4f**). Together, these results suggest that H-NS and SspA are epistatic for CRISPR interference, with H-NS acting downstream of SspA on the regulation of CRISPR interference-mediated anti-phage defence.

### SspA regulates CRISPR adaptation independently of H-NS

Given that SspA may regulate CRISPR interference via H-NS, we sought to determine whether SspA, in turn, regulates CRISPR adaptation via H-NS as well. To do so, we performed deep sequencing of the CRISPR arrays from samples harvested 3h post α_vir_ infection in liquid cultures of wild-type, Δ*hns*, Δ*sspA*, and Δ*hns* Δ*sspA* mutants harbouring either T or NT plasmids. We found no differences in the rates of CRISPR adaptation across conditions, except in the Δ*hns* + T cultures, which substantially increased rates of CRISPR adaptation (**Fig 5a**). These new spacers were primarily α_vir_ derived (**Fig 5b**), and that the majority of the acquired spacers are found immediately downstream and on same strand as the immunising spacer, consistent with primed CRISPR adaptation (**Fig 5c-d, Extended Data Figure 5a**). Interestingly, we saw a substantial decrease in α_vir_ derived spacers in the Δ*hns* Δ*sspA* + T conditions (**Extended Data Figure 5c**). Although the rates of CRISPR adaptation in the Δ*hns* + T condition were low (0.5% of CRISPR arrays expanded, i.e., 5 cells per thousand with a newly expanded array) and could not explain the defence demonstrated by the Δ*hns* + T cultures at the time of sample collection (**Fig 4f**), our results underscore the requirement for SspA for adequate primed CRISPR acquisition, in a closer-to-natural and defence-relevant setting.

**Figure 5.**
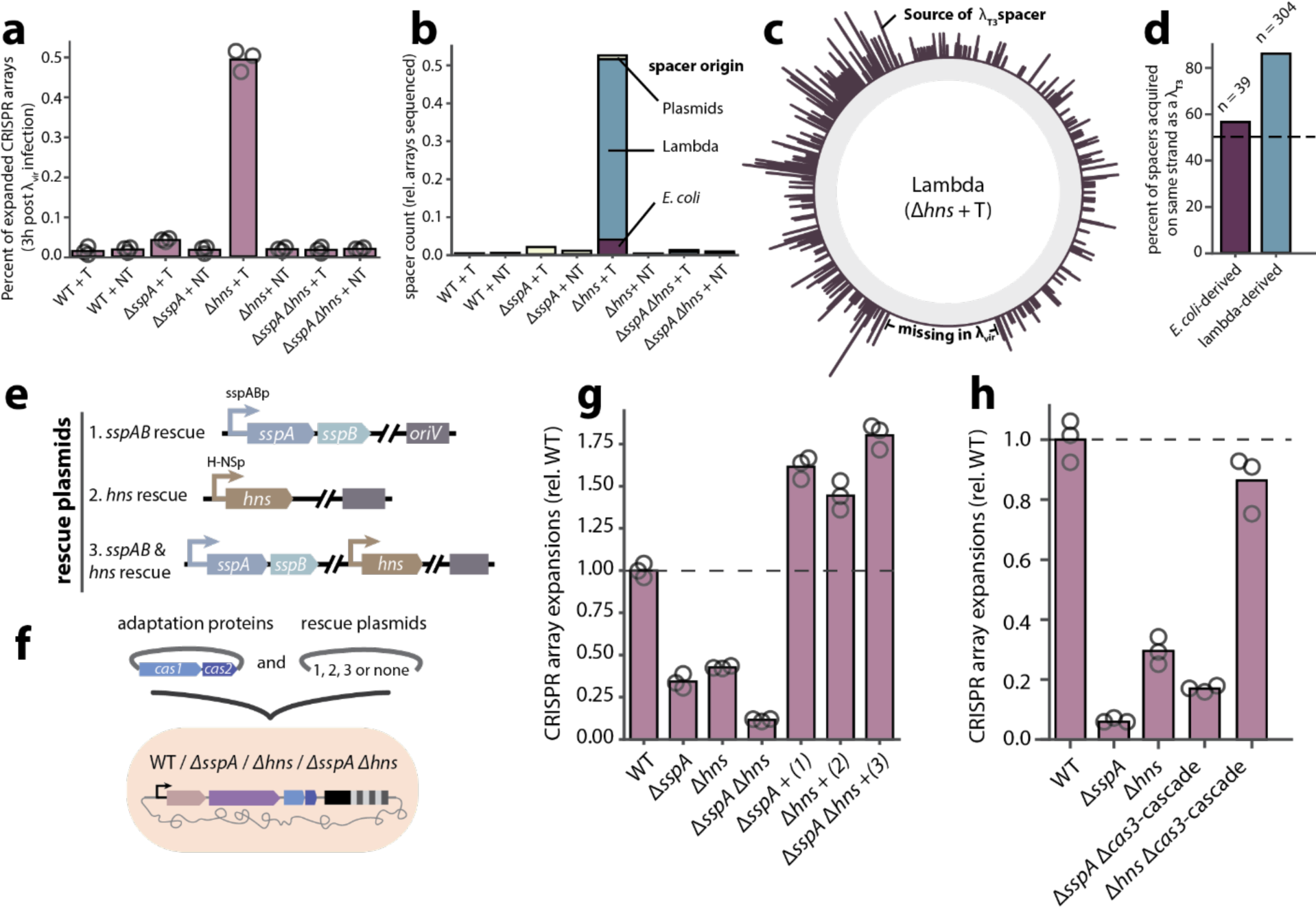
SspA regulates CRISPR adaptation independently of H-NS. **a**. Deep-sequencing based measurement of the rates of new spacer acquisition in strains pre-immunised with either a T or NT defence plasmid, harvested 3h post λvir infection in liquid culture and growth at 30°C. Open circles represent biological replicates (*n* ≥ 3), bars are the mean (one-way ANOVA effect of strain *P*< <0.0001; Sidak’s corrected multiple comparisons for wild-type +T vs. knockouts +T, Δ*sspA* P=082553, Δ*hns* P<0.0001, Δ*sspA* Δ*hns* P=0.999999; Δ*sspA* +T vs. knockouts *+*T, Δ*hns* P<0.0001, Δ*sspA* Δ*hns* P=0.154762; Δ*hns +*T vs. Δ*hns* +NT P<0.0001; Δ*hns +*T vs. Δ*sspA* Δ*hns* +T P<0.0001). **b.** Breakdown of normalised spacer count (total number of new spacers / number of CRISPR arrays sequenced) according to spacer origin (*E. coli*, lambda or plasmid) and strain of interest. **c.** Binned coverage plot of Δ*hns* + T newly acquired spacers across the lambda genome (outer, purple). The location of the T immunisation spacer is shown on the lambda genome; “missing in /\vir” indicates a genomic region missing in our strain of /\vir. **d**. Percent of spacers acquired that are on the same strand as the T immunisation spacer, according to the spacer source (*E. coli* or lambda). **e**. Schematic of the *sspAB* and *hns* operonic rescue plasmids. All plasmids are low (∼5) copy, and encode either 1. The sspAB operon, 2. The hns operon, or 3. both, under their native regulation. **f**. Schematic of the CRISPR adaptation assays in wild-type, *sspA* and/or *hns* mutant strains. Strains were electroporated with pSCL565 and rescue plasmids 1., 2., or 3. (see **e**.), and assessed for their ability to acquire new spacers into the endogenous CRISPR I array. **g**. PCR-based detection of new spacer acquisition into the CRISPR I array of wild-type, of WT, Δ*sspA*::*FRT,* Δ*hns*::*FRT* and Δ*sspA*::*FRT* Δ*hns*::*FRT* strains harbouring pSCL565 and rescue plasmids 1., 2., or 3. (see **e**.), after growth for 48h in liquid culture. Open circles represent biological replicates (*n* ≥ 3), bars are the mean. Horizontal dashed line represents the mean rate of spacer acquisition in the wild-type strain (one-way ANOVA effect of strain *P*< <0.0001; Sidak’s corrected multiple comparisons for wild-type vs. knockouts, Δ*sspA* P<0.0001, Δ*hns* P<0.0001, Δ*sspA* Δ*hns* P<0.0001; Δ*sspA* vs. knockouts, Δ*hns* P=0.714182, Δ*sspA* Δ*hns* P=0.002269, Δ*sspA + sspAB* rescue P<0.0001; Δ*hns* vs. knockouts, Δ*sspA* Δ*hns* P<0.0001, Δ*hns + hns* rescue P<0.0001; Δ*sspA* Δ*hns* vs. Δ*sspA* Δ*hns* + *sspA* & *hns* rescues P<0.0001). **h**. PCR-based detection of new spacer acquisition into the CRISPR I array of WT, Δ*sspA*::*FRT,* Δ*hns*::*FRT*, Δ*sspA*::*FRT Δcas3-Cascade::Cm^R^* or Δ*hns*::*FRT Δcas3-Cascade::Cm^R^* strains harbouring pSCL565 after growth for 48h in liquid culture. Open circles represent biological replicates (*n* ≥ 3), bars are the mean (one-way ANOVA effect of strain *P*<0.0001; Sidak’s corrected multiple comparisons for wild-type vs. knockouts, Δ*sspA* P<0.0001, Δ*hns* P<0.0001, Δ*sspA* Δ*cas3-cascade* P<0.0001, Δ*hns* Δ*cas3-cascade* P=0.125466; Δ*sspA* vs. Δ*hns* P=0.004161; Δ*sspA* vs. Δ*sspA* Δ*cas3*-*cascade* P=0.310715; Δ*hns* vs. Δ*hns* Δ*cas3-cascade* P<0.0001; Δ*sspA* Δ*cas3*-*cascade* vs. Δ*hns* Δ*cas3*-*cascade* P<0.0001). Horizontal dashed line represents the mean rate of spacer acquisition in the wild-type strain. Additional statistical details in **Supplemental Table 1**.

We next sought to determine whether SspA modulates naïve CRISPR adaptation via H-NS. For this, we used the Δ*sspA::FRT*, Δ*hns::FRT* and Δ*sspA::FRT* Δ*hns::FRT E. coli* strains (**Fig 5e**), and assessed the mutants’ ability to acquire new spacers after co-electroporation of pSCL565 alongside a low (∼5) copy rescue plasmid encoding the *sspAB* operon, *hns* operon, or both, under their native genomic contexts and regulation (**Fig 5f**). We found that Δ*sspA*, Δ*hns*, and Δ*sspA* Δ*hns* mutant strains all showed defects in CRISPR adaptation, with the double Δ*sspA* Δ*hns* mutant showing the strongest defect (**Fig 5g**). Complementation of the knockout strains with their respective rescue plasmids restored CRISPR adaptation to levels comparable to wild-type.

Since H-NS deletion de-represses CRISPR interference (Fig **4d-f**), we hypothesised that its effect on CRISPR adaptation could be indirect, through the removal of cells that acquired genome-derived spacers via CRISPR interference-mediated self-targeting. To remove the confounding effect of increased self-targeting in the Δ*hns* background, we built Δ*cas3-cascade::cm^R^* knockouts on top of the Δ*sspA* and Δ*hns* genetic backgrounds, and assessed the mutants’ ability to acquire new spacers after electroporation with pSCL565. We found that although the Δ*sspA* Δ*cas3-cascade::cm^R^* mutant still remained substantially CRISPR adaptation deficient, the Δ*hns* Δ*cas3-cascade::cm^R^* mutant recovered CRISPR adaptation to levels comparable to wild-type (**Fig 5h**). This confirmed that the apparent CRISPR adaptation deficiency of the Δ*hns* mutant was caused by self-targeting through de-repression of CRISPR interference, and not additional effects on CRISPR adaptation. Taken together, our data supports a role for SspA in CRISPR adaptation that is independent of H-NS.

## DISCUSSION

We developed a novel negative selection CRISPRi screen, designed around the concept of stimulated CRISPR self-immunity, to identify potential host factors that participate in CRISPR adaptation in *E. coli*. We identified a new host factor in our screen, SspA. In validation experiments, adaptation assays and downstream analysis of newly acquired spacers revealed that a *sspA* knockout mutant is consistently and significantly defective in naïve CRISPR adaptation, despite no other noticeable differences in the features of its acquired spacers when compared to the wild-type parental strain. Further, we found that mutations that abolish SspA’s ability to bind to the RNA Polymerase complex cause a loss-of-adaptation phenotype, suggesting that SspA acts as a transcriptional-level regulator of CRISPR adaptation. A series of phage sensitivity and CRISPR adaptation assays revealed that SspA regulates CRISPR adaptation independently of H-NS, a known regulator of CRISPR interference-mediated anti-phage defence and a member of the SspA regulon. Taken together, our data support independent control of CRISPR adaptation and interference downstream of SspA.

We find that our data is consistent with a model where the immunisation and interference steps could occur separately, perhaps even temporally so. We speculate that phage infection could trigger the rapid accumulation of SspA^67^, opening a window for the acquisition of new spacers; this window may close rapidly as the levels of SspA decline, but this sudden SspA accumulation may be enough to cause downregulation of H-NS^60^, thus opening a second window for CRISPR interference to occur (**Fig 6**). However, more studies are required to determine whether the sudden accumulation of SspA in response to phage infection is a ubiquitous response beyond lambda, what phage element or phage-induced signal triggers this sudden spike in SspA levels, and whether this spike is indeed sufficient to significantly deplete levels of H-NS and open a window for CRISPR interference to occur. Further, though our data strongly suggests that SspA acts on CRISPR adaptation at a transcriptional level, additional work is needed to discover the target(s) of the SspA-mediated transcriptional rewiring.

**Figure 6.**
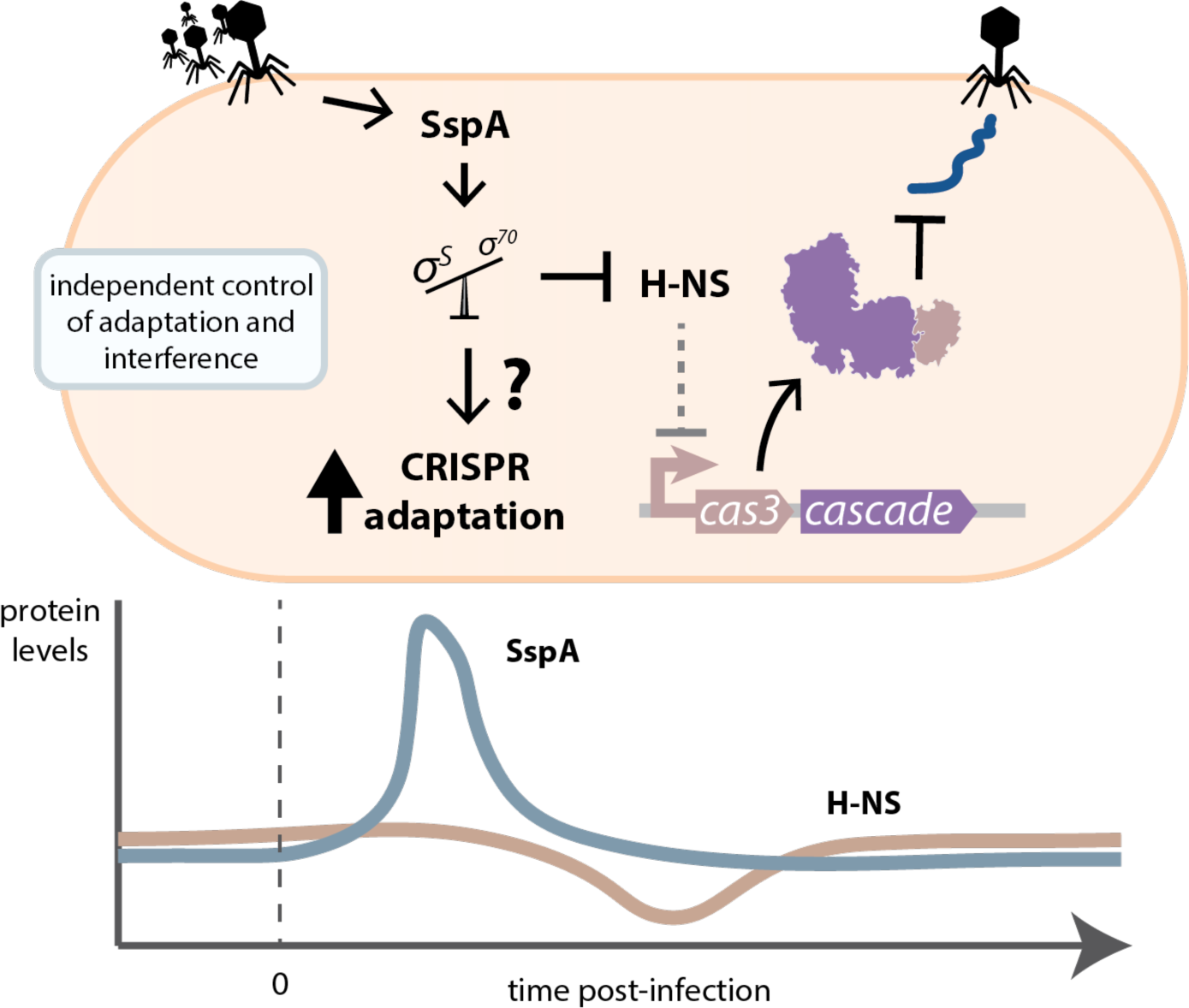
Proposed model for the independent control of CRISPR adaptation and interference. In both cases, the regulation of CRISPR immunity happens downstream of SspA, through its role as a global transcriptional rewiring agent.

Though our screen revealed SspA as a novel regulator of CRISPR adaptation, we did not identify host factors involved in the repair of the CRISPR array. Although *polA* was a promising hit, with *in vitro* evidence that its Klenow fragment is capable of repairing CRISPR arrays that have been cleared of the Cas1-Cas2 integrases^22^, our results do not support this role. Though we found that CRISPR adaptation levels were significantly diminished in the *polA* ΔKlenow mutant, this decrease was attributable to the loss of acquisition of plasmid-derived spacers; The loss-of-adaptation phenotype seen in the Δ*pcnB* mutant is likely due to a similar effect, as *pcnB* has been shown to be required for copy number maintenance of ColE1 and other plasmids^72^. We cannot, however, rule out a role for *polA* in array repair, though it is conceivable that there is redundancy in host factors capable of this task. Indeed, functional redundancy of host factors is a possible explanation for not capturing the comprehensive set of these proteins. We anticipate that more complex combinatorial knockdown and activation screens could be used to tackle this problem. Furthermore, we believe that pairing genetic screens such as our CRISPRi screen with orthogonal physical screens, such as proximity labelling and pull-down assays^73^, will yield rich and informative datasets, which are likely to uncover a more comprehensive set of host factors required for CRISPR adaptation.

## METHODS

### Bacterial strains and culturing

All strains used in this study can be found in Supplementary Table 2. Wild-type *E. coli* K-12 W3110 (BW25113) strain, generously provided by Joseph Bondy-Denomy, was used for all experiments in this study, unless specified. *E. coli* K-12 MG1655 and LC-E75^43^ (derivative of MG1655, Addgene #115925) were used for the CRISPRi screen. *E. coli* NEB-5-alpha (NEB C2987) was used for plasmid cloning. Keio collection^46^ single-gene knock-out (KO) mutants, derivatives of BW25113, were generously provided by Carol Gross.

Additional deletions on Keio single-gene KO backgrounds were generated by λRed recombinase-mediated insertion of an FRT-flanked chloramphenicol (Cm^R^) resistance cassette^74^. This cassette was amplified from pKD3^74^ (Addgene #45604) with homology arms (50bp each) corresponding to the genomic sequences immediately up- and downstream of the intended deletion site. This amplicon was electroporated into the Keio strains expressing the λRed recombinase from pKD46^74^. Clones were isolated by selection on LB + chloramphenicol (10 µg/mL) plates. After PCR genotyping and sequencing to confirm locus-specific insertion, the chloramphenicol and pre-existing kanamycin cassettes was excised by transient expression of FLP recombinase from pE-FLP^75^ (Addgene #45978) to leave a single FRT scar, whenever specified in the text (i.e., Δ*gene*::*FRT*).

The *polA* ΔKlenow mutant was generated by λRed recombinase-mediated insertion of an FRT-flanked Cm^R^ resistance cassette into the Klenow fragment of *E. coli* BW25113 Polymerase I. This cassette was amplified from pKD3 with homology arms (50bp each), corresponding to the genomic regions flanking the Klenow fragment, as reported previously^47^. This amplicon was electroporated into BW25113 expressing the λRed recombinase from pKD46. Clones were isolated by selection on LB + chloramphenicol (10 µg/mL) plates. PCR genotyping and sequencing confirmed the locus-specific insertion.

For the CRISPRi screen and CRISPR-Cas adaptation experiments, LB containing 1.5% w:v agar was used to grow strains on plates (growth at 37°C until single colonies became visible, usually ∼16h). Strains were subsequently grown in LB broth at 37°C with 250 r.p.m. shaking, with appropriate inducers and antibiotics as described below.

For CRISPR-Cas defence experiments, strains were grown in LB broth supplemented with 10 mM MgSO4 and 0.2% maltose at 30°C with 250 r.p.m. shaking, with appropriate inducers and antibiotics as described below. For plaque assays, cells were mixed with top agar (0.5% w:v LB agar, supplemented with 10 mM MgSO4 and 0.2% maltose and the appropriate antibiotics) poured over LB plates supplemented with the appropriate antibiotics, and grown at 30°C overnight.

Inducers and antibiotics were used at the following working concentrations: 2 mg/mL L-Arabinose (GoldBio A-300), 1 mM IPTG (GoldBio I2481C), 1mM m-Toluic acid, 1 ug/mL anhydrotetracycline, 35 µg/mL kanamycin (GoldBio K-120), 25 µg/mL spectinomycin (GoldBio S-140), 100 µg/mL carbenicillin (GoldBio C-103), 25 µg/mL chloramphenicol (GoldBio C-105).

### Phage strains and culturing

A virulent variant of phage Lambda (αvir)^76^, generously provided by Luciano Marraffini, was used throughout this study. αvir was propagated on BW25113 grown in LB at 30°C, based on previous studies^77^. Briefly, overnights of *E. coli* BW25113 were grown at 30°C in 5mL LB + 10 mM MgSO4 and 0.2% maltose. The next day, 300uL of bacterial culture was infected with 10uL of serial dilutions of αvir in LB + 10 mM MgSO4 and 0.2% maltose, incubated at 30°C for 15min, and added to 5mL top agar, mixed gently and poured over LB agar plates. Plates were grown overnight at 30°C. Plates from the dilution series that showed evidence of confluent lysing of *E. coli* were covered in 5mL LB supplemented with 10 mM MgSO4 and 0.2% maltose, placed on a shaker to agitate gently at room temperature for 2h. Then, the lysate was transferred to a 15mL conical tube, centrifuged at 4500g x 15min to remove the bacterial debris, and filtered through a 0.2um filter. Phage titres were determined by preparing 1:10 dilutions of αvir in LB supplemented with 10 mM MgSO4 and 0.2% maltose, and spotting 2.5uL of the dilutions over top agar lawns of BW25113, which had been previously prepared by mixing 100uL of the overnight culture with 5mL of top agar (0.5% w:v LB agar, supplemented with 10 mM MgSO4 and 0.2% maltose) and poured over LB agar plates. Serial dilutions of αvir were prepared in LB supplemented with 10 mM MgSO4 and 0.2% maltose, and 2.5uL of each dilution was spotted on the top agar using a multichannel pipette. Plates were tilted to allow phage spots to drip down the plate for easier quantification, and left to dry completely at room temperature. Plates were incubated at 30°C overnight.

### Plasmids

Plasmid pSCL565, encoding an IPTG-inducible *E. coli* Cas1-Cas2 cassette, spectinomycin resistance cassette and a pCDF ori, was constructed by PCR amplification of pCas1+2^14^ (Addgene #72676) to replace the T7 promoter by an IPTG-inducible Lac promoter.

Plasmid pSCL563, encoding an m-Tol-inducible *E. coli* Cas3-Cascade operon, carbenicillin resistance cassette and a pRSF ori was constructed by Gibson cloning.

The *sspAB* rescue set of plasmids, designed to rescue the loss of CRISPR adaptation phenotype of the *ΔsspA* mutant, were constructed by first Gibson cloning the *sspAB* operon (including 236bp upstream of *sspA,* containing the predicted promoter^78^ between *rpsI* and *sspA*) into a low copy plasmid backbone (pSC101 ori) containing a carbenicillin resistance cassette. This yielded pSCL735 (*sspAB* rescue). Variants of the *sspAB* operon were generated by targeted PCRs to yield pSCL747 (*sspA* rescue, not tested because toxic in Δ*sspA* background); pSCL748 (*sspB* rescue); pSCL751 (*sspA* frameshifted AN^5–6^>AQ^5–6^ GCC|AAC>GC**T**|*CAA*|C + *sspB*); pSCL752 (*sspA + sspB* frameshifted PR^9–10^>PS^9–10^ CCA|CGT>CCA|***T****CG*|T); pSCL753 (*sspA* frameshifted AN^5–6^>AQ^5–6^ GCC|AAC>GC**T**|*CAA*|C + *sspB* frameshifted PR^9–10^>PS^9–10^ CCA|CGT>CCA|***T****CG*|T); and pSCL770 (*sspA* PHP^84–86^>AAA^84–86^ + *sspB*).

The CRISPR defence set of plasmids, designed to pre-immunise *E. coli* strains against αvir by expressing an *E. coli* CRISPR-I array with a first spacer encoding either a Target (complementary to the α genome^1,70^ or a Non-Target (NT) spacer, were constructed by cloning a spacer1-swapped *E. coli* CRISPR-I array (Cas-adjacent array in K-12 *E. coli*) into a high copy plasmid backbone (ColE1) containing a kanamycin resistance cassette. This yielded pSCL787 (Target, spacer1 complementary to the αvir *R* gene^1^ and pSCL788 (Non-Target, spacer1 complementary to the *S. cerevisiae ade2* gene).

The *sspAB*-*hns* rescue set of plasmids, designed to rescue the loss of CRISPR adaptation phenotype of the Δ*hns* and *ΔsspA* Δ*hns* mutants, were constructed by Gibson cloning the *hns* operon (including 419bp upstream and 122bp downstream of *hns,* containing the predicted promoter^78^, regulatory and terminator regions contained between *tdk*-*hns-galU*, respectively) and/or *sspAB* operons (as above) into a low copy plasmid backbone (pSC101 ori) containing a carbenicillin resistance cassette. This yielded pSCL785 (*hns* rescue) and pSCL786 (*hns // sspAB* rescue).

pSCL832 was constructed from pSCL565 by swapping the *E. coli* Cas1-Cas2 CDS with an eGFP CDS via Gibson Assembly.

Additional plasmid information can be found in Supplementary Table 3.

### CRISPRi adaptation host factor screen

LC-E75^43^, a derivative of MG1655 *E. coli* encoding a Tetracycline-inducible dCas9 cassette integrated at the Phage 186 *attB* site, and *E. coli* MG1655 were electroporated with pSCL565, and transformants were isolated on LB + spectinomycin after overnight growth at 37°C. Single colonies were inoculated into 5mL LB + spectinomycin, and grown overnight. Each experiment was repeated 3 times in triplicates, for a total of 9 paired LC-E75 (experiment) – MG1655 (control) screens.

The next day, cultures were electroporated with a library of 92,919 sgRNAs (psgRNA^43^ Pooled Library #115927, Addgene), targeting coding and non-coding regions across the *E. coli* genome, as described in^43,44^. Briefly, 4mL of the overnight cultures were diluted into 400mL LB + spectinomycin, and grown for 2h at 37°C with shaking (250 r.p.m.). Cells were then subjected to an electroporation prep: cultures were split into 50mL falcon tubes, chilled on ice for 10min, and pelleted at 4000g for 15min at 4°C. The supernatants were discarded, and cells were washed with 30mL of ice-cold ultra-pure, DNAse/RNAse free, pyrogen free H2O (updH2O). The resuspended cultures were chilled on ice for another 10min, then pelleted at 4000g for 15 min at 4°C. These wash steps were repeated twice, for 3 total washes. After the last wash, cells were resuspended in 600uL of 10% glycerol in updH2O (∼800uL final volume).

Then, 180uL of cells were added to 0.2cm gap electroporation cuvettes (BioRad #1652086), and ∼1ug of the sgRNA library was mixed with the cells (total volume in electroporation cuvette < 200uL). Cells were electroporated with the following settings: 2.5 kV, 25uF, 200Ω. After the pulse, cells were quickly recovered in 25mL of pre-warmed LB + spectinomycin, and placed in a shaking incubator for 1h at 37°C. The cultures were then transferred into 75mL of pre-warmed LB + spectinomycin + kanamycin, and dilutions were plated on LB + spectinomycin + kanamycin to estimate CFUs.

The next day, CFUs were estimated, and the experiments were continued only if the library coverage was estimated to be >1000x. If so, 20mL of the overnight cultures were diluted in 1L warmed LB + spectinomycin + kanamycin + 1uM anhydrotetracycline (aTc); the remainder of the overnight cultures was collected by centrifugation for pre-experiment library quantification.

Cultures were grown for 3h, after which the electroporation prep was performed as described above. After the last centrifugation step, each pellet was resuspended in 150uL of a mix of *murA* targeting pre-spacer oligonucleotides and ∼1ug of pSCL563 in updH2O. The *murA* targeting prespacer mix was prepared by combining and annealing complementary single-stranded oligos that encode a prespacer targeting the essential gene *murA* (F and R sequences: AGGTTATGGCAACCGATCTGCGTGCATCAGCAAGC;

GCTTGCTGATGCACGCAGATCGGTTGCCATAACCT), to a final concentration of 3.125 uM per oligo. After electroporation, cells were rescued with 5mL of pre-warmed LB + carbenicillin + kanamycin + 1 mM m-Toluic acid + 1uM aTc, and placed in a shaking incubator for 1h at 37°C. Then, these cultures were then transferred into 20 mL of pre-warmed LB + carbenicillin + kanamycin + 1 mM m-Toluic acid + 1uM aTc, and placed in a shaking incubator overnight at 37°C. Cultures (post-experiment library samples) were harvested the next day by centrifugation, 4000g x 30min, followed by plasmid extraction using the Qiagen Plasmid Plus Midi kit (cat. no. 12143).

Sequencing of the sgRNA libraries was performed as follows. 1uL of the plasmid extractions were used as template in 50uL PCR reactions, using 37uL of updH2O, 10uL 5X Q5 reaction buffer, 1uL 10mM dNTPs, 1uL Q5 Hot Start HiFi DNA polymerase and 0.25uL 100uM Forward and Reverse primers. The primers used contained Illumina adapters to make the amplicons compatible with our downstream sequencing prep, as well as 1-5 random nucleotides between the Illumina adapter and the annealing sequence to introduce diversity into the sequencing library. The PCR reaction was run using the standard recommended Q5 cycling conditions: 98°C initial denaturation x 30s; 30 cycles of 98°C x 10s, 62°C x 30s, 72°C x 30s; final extension of 2min at 72°C. Amplicons were then cleaned up using AMPure XP beads (A63880), indexed using custom indexing oligos, and sequenced on an Illumina NextSeq instrument with ∼2million reads per biological replicate. A list of primers can be found in Supplementary Table 4.

### Fluorescence-based monitoring of the Lac promoter activity

*E. coli* BW25113 (control) and Δ*sspA* strains were transformed with pSCL832 by electroporation, and transformants were isolated on LB + spectinomycin after overnight growth at 37°C. Single colonies (*n* ≥ 3) were inoculated into 3mL of LB + spectinomycin and grown overnight with 250 r.p.m shaking at 37°C. The next day, cultures were diluted 1:100 in 3mL of LB + spectinomycin and grown to log phase (∼4h). Subsequently, OD600 of the cultures was measured on a Spectramax i3 plate reader, and cultures were normalised to an OD600 = 0.05. 200uL of cultures were placed on clear-bottom plate and incubated at 37°C on a Spectramax i3 plate reader, with fluorescence readings (wavelength = 508nm) every 30s for a total of 7.5h.

### qPCR

*E. coli* BW25113 (control) and Δ*sspA* strains were transformed with pSCL565 by electroporation, and transformants were isolated on LB + spectinomycin after overnight growth at 37°C. Single colonies (*n* ≥ 3) were inoculated into 3mL of LB + spectinomycin and grown overnight with 250 r.p.m shaking at 37°C. The next day, cultures were diluted 1:100 in 3mL of LB + spectinomycin and grown to log phase (∼4h). Then, 1mL of cultures was harvested by centrifugation (21,000 g x 1min), then resuspended in 250uL of updH2O. These samples were heated to 95°C for 15min, then placed on ice to cool. Then, lysates were treated with 2 units of Proteinase K (NEB) for 30min, followed by Proteinase K inactivation by incubation at 95°C for 10min. Lastly, lysates were centrifuged at 21,000 g for 2min, and supernatants were diluted 1:500 in updH2O. 5uL of the diluted supernatant was used in 20uL qPCR reactions, set up using the NEB Luna Universal qPCR Master Mix following the manufacturer’s instructions. qPCR Primers were designed to target pSCL565’s CDF ori and *cas1* regions, using the genomic *ompA* as a reference. Primers are listed in Supplementary Table 4.

### CRISPR-Cas adaptation experiments Naïve CRISPR-Cas adaptation

*E. coli* BW25113 (control) and strains of interest were transformed with pSCL565 by electroporation, and transformants were isolated on LB + spectinomycin after overnight growth at 37°C. In the case of “plasmid rescue” experiments, strains of interest we co-transformed with pSCL565 and the rescue plasmid by electroporation, and transformants were isolated on LB + spectinomycin + carbenicillin after overnight growth at 37°C.

Single colonies (*n* ≥ 3) were inoculated into individual wells of a 96-well deep well plate containing 500uL of LB + spectinomycin (and carbenicillin, if needed), and grown for 48h with 1000 r.p.m shaking at 37°C. After 48h of growth, 75uL of the cultures were mixed with 75uL of updH2O, heated to 95°C for 10min, and spun-down. 0.5uL of the supernatant was used as template for 25uL PCR reactions (same recipe and cycling protocol as above). We designed primers to amplify a region of the *E. coli* CRISPR-II array, contained between the end of the Leader sequence and the second pre-existing spacer. To reduce the number of indices needed per sample, we designed 3 barcoded F primers (one per biological replicate) to amplify the CRISPR arrays – these would enable us to pool the samples post-CRISPR array amplification, and de-multiplex the biological replicates during data analysis. A list of primers can be found in Supplementary Table 4.

In some cases, CRISPR array expansions are visible on an agarose gel as laddering caused by larger arrays (expanded) migrating slower than the shorter parental arrays. We visualised this by running 5uL of the pooled PCR products on Invitrogen 2% Agarose SYBR safe E-Gels (A42135). Gels were re-stained with SYBR Gold before imaging.

### CRISPR-Cas interference experiments Phage plaque assays

BW25113 (control), Δ*sspA*::FRT, Δ*hns*::FRT, and Δ*sspA*::FRT Δ*hns*::FRT strains were transformed with plasmids encoding either Target or Non-Target CRISPR-I arrays (pSCL787 and pSCL788, respectively), and transformants were isolated on LB + kanamycin after overnight growth at 37°C. Single colonies (*n* ≥ 3) were inoculated into 3mL of LB + kanamycin supplemented with 10 mM MgSO4 and 0.2% maltose, and grown overnight with 250 r.p.m shaking at 30°C. The next day, top agar lawns of each bacterial culture were prepared by mixing 100uL of overnight cultures with 5mL of top agar (0.5% w:v LB agar, supplemented with 10 mM MgSO4 and 0.2% maltose and kanamycin). Top agar mixtures were poured over LB agar + kanamycin plates and left to dry at room temperature, partially open by a sterilizing flame. Serial dilutions of λvir were prepared in LB supplemented with 10 mM MgSO4 and 0.2% maltose, and 2.5uL of each dilution was spotted on the top agar using a multichannel pipette, and left to dry completely at room temperature. Plates were incubated at 30°C overnight.

Efficiency of plating was calculated as the number of plaques formed by λvir on lawns of a strain harbouring pSCL787 (Target) divided by the plaques formed by λvir on lawns of a strain harbouring pSCL788 (Non-Target). Full plaque assay plates for all *n* = 3 biological replicates in Extended Data Fig 5.

### Phage resistance infection growth curves

BW25113 (control), Δ*sspA*::FRT, Δ*hns*::FRT, and Δ*sspA*::FRT Δ*hns*::FRT strains were transformed with plasmids encoding either Target or Non-Target CRISPR-I arrays (pSCL787 and pSCL788, respectively), and transformants were isolated on LB + kanamycin after overnight growth at 37°C. Single colonies (*n* ≥ 3) were inoculated into 3mL of LB + kanamycin supplemented with 10 mM MgSO4 and 0.2% maltose, and grown overnight with 250 r.p.m shaking at 30°C. The next day, cultures were diluted 1:100 in 3mL of LB + kanamycin supplemented with 10 mM MgSO4 and 0.2% maltose and grown to log phase (∼4h). Subsequently, OD600 of the cultures was measured on a Spectramax i3 plate reader, and cultures were normalised to an OD600 = 0.05. 200uL of cultures was infected with a range of MOIs (10 σ 10^-8^), using serial dilutions of λvir prepared in LB supplemented with 10 mM MgSO4 and 0.2% maltose. Cultures were loaded on clear-bottom plate and incubated at 30°C on a Spectramax i3 plate reader, with OD600 readings every 2.5mins for a total of 16h.

### CRISPR-Cas primed adaptation after phage infection

BW25113 (control), Δ*sspA*::FRT, Δ*hns*::FRT, and Δ*sspA*::FRT Δ*hns*::FRT strains were transformed with plasmids encoding either Target or Non-Target CRISPR-I arrays (pSCL787 and pSCL788, respectively), and transformants were isolated on LB + kanamycin after overnight growth at 37°C. Single colonies (*n* ≥ 3) were inoculated into 3mL of LB + kanamycin supplemented with 10 mM MgSO4 and 0.2% maltose, and grown overnight with 250 r.p.m shaking at 30°C. The next day, cultures were diluted 1:100 in 3mL of LB + kanamycin supplemented with 10 mM MgSO4 and 0.2% maltose and grown to log phase (∼4h). Subsequently, OD600 of the cultures was measured on a Spectramax i3 plate reader, and cultures were normalised to an OD600 = 0.05. 200uL of cultures was infected with αvir at an MOI of 0.1, and cultures were loaded on clear-bottom plate and incubated at 30°C on a Spectramax i3 plate reader, with OD600 readings every 1.5mins for a total of 3h.

After 3h, the cultures were harvested by centrifugation, resuspended in 100uL of updH2O, heated to 95°C for 10min, and spun-down. 1uL of the supernatant was used as template for 25uL PCR reactions (same recipe and cycling protocol as above). We designed primers to amplify a region of the *E. coli* CRISPR-II array, contained between the end of the Leader sequence and the second pre-existing spacer. The barcoded F primer approach, described above, was used to pool PCRs and de-multiplex biological replicates during data analysis.

### Protein model structures

Protein model coordinates were retrieved from the RSCB Protein Data Bank (codes 7DY6 and 8ET3). Figures were prepared using UCSF ChimeraX^79^.

### Data analysis

The data analysis for this project can be broken down into 5 modules: (1) processing of the sequencing reads to extract, count, and group sgRNAs by gene/gene-adjacent regions; (2) generate binned coverage plots of sgRNAs across the *E. coli* genome; (3) identify the statistically enriched/depleted sgRNAs, using PyDESeq2^45^, a Python implementation of DEseq2^80^; (4) quantify the rates of CRISPR adaptation; and (5) extract new spacers perform spacer analysis. All data analysis was performed in Jupyter Lab^81^, and all code to replicate this analysis can be found here: https://github.com/Shipman-Lab/CRISPRi_host_factor_screen.

### Sequencing data processing: from reads to sgRNA counts

First, fastq reads were trimmed using sickle-trim^82^. For each fastq, a counter of sgRNAs was generated by extracting the sgRNA from each read, provided that this sgRNA could be found in the original synthesised psgRNA library^43^. Then, the sgRNAs were BLASTed^83^ against the *E. coli* MG1655 genome and the top hit was saved. For each sample, a DataFrame of genomic_location–sgRNA–count was generated, and used for downstream analysis. All data corresponding to the screens can be found in Supplementary Tables 5-6. The Jupyter Notebook for this analysis can be found here: https://github.com/Shipman-Lab/CRISPRi_host_factor_screen/blob/main/blast_screen_hits_clean.ipynb

### Binned coverage plot of sgRNAs across the *E. coli* genome

We generated occupancy arrays for each sample, using counts generated above. These arrays contain cumulative counts of sgRNAs per base, i.e., occupancy *O* = [c1, c2, …, c*n*], where *n* is the size of the *E. coli* MG1655 genome and ci are the total sgRNA counts at that position. We then normalised the counts to the total sgRNA count in that sample, i.e., *O* = [c1/sum_sgRNAs, c2/sum_sgRNAs, …, c*n*/sum_sgRNAs], where sum_sgRNAs is total sgRNA count. Next, we calculated the mean occupancy for the experimental and control conditions, i.e., *OLC-E75* = (Obiorep1 + Obiorep2 + … +Obiorep9) / 9, where *OLC-E75* is the mean occupancy for the experimental condition, and Obiorep_i are the normalised counts for each biological replicate of the screen run in the experimental condition. Lastly, we calculated the delta occupancy, or difference between the mean sgRNA occupancies of the experimental and control conditions, and posteriorly calculated the mean delta sgRNA occupancy in a sliding window, in the interest of interpretability. We used pyCirclize^84^ to generate the final occupancy plot.

The Jupyter Notebook for this analysis can be found here: https://github.com/Shipman-Lab/CRISPRi_host_factor_screen/blob/main/plot_genome_coverage_clean.ipynb

### Identification of enriched/depleted sgRNAs

We performed statistical testing for enriched/depleted sgRNAs from binned sgRNA (sum of all sgRNAs per gene) count data generated in (1) using the PyDESeq2 package^45^, and compared each experimental sample to its paired control, and controlled for pre-experimental variation in the relative sgRNA library composition by including the sgRNA counts from the pre-experiment library as an interactor factor (i.e., sgRNA_counts ∼ input_lib_counts + knockdown (yes/no)). Genes that have less than a total of 10 reads for all of their sgRNAs in the dataset were removed from the analysis. The log2FoldChange (log2FC) value represents the enrichment or depletion of each gene. The lists of all genes, log2FC and adjusted p-values can be found in Supplementary Table 6. The Jupyter Notebook for this analysis can be found here: https://github.com/Shipman-Lab/CRISPRi_host_factor_screen/blob/main/deseq2_volcano_allhits_clean.ipynb

### Quantification of the rates of CRISPR adaptation

First, fastq reads were trimmed using sickle-trim. For each fastq, we filtered for reads containing the Leader-repeat junction of the *E. coli* CRISPR-II array. We then identified newly acquired spacers from the array sequences by recursive identification of CRISPR repeats and comparison of putative new spacers to pre-existing spacers in the array, using a lenient search algorithm allowing for a maximum of 3bp mismatches. We generated sums of new expansions in CRISPR arrays per condition, and used these to calculate the rate of CRISPR adaptation (100 * number of newly expanded CRISPR arrays / total number of arrays sequenced). Lastly, we normalised the rate of CRISPR adaptation for each condition by the wild-type rate CRISPR adaptation, so as to make inter-experiment comparisons feasible and more interpretable. All normalised rates corresponding to the CRISPR adaptation experiments, as well as the “run” label (i.e., batch in which the experiment were run and sequenced) can be found in Supplementary Tables 7. The Jupyter Notebook for this analysis can be found here: https://github.com/Shipman-Lab/CRISPRi_host_factor_screen/blob/main/spacer_fishing_clean.ipynb

### Newly-acquired spacer analysis

Spacer analysis involves in several steps:

#### Extraction of new spacers

We began by using the same recursive new spacer search algorithm described above to extract new spacers. In parallel to extracting spacers, we also stored information regarding the total number of arrays sequenced and the fraction of those that were expanded, to use as normalisation for comparisons across samples that might vary in sequencing depth or quality.

#### Identification of spacer origin

Next, we generated a counter of newly acquired spacers and their frequencies. We used this to generate FASTA files of new spacers and their counts, which were subsequently BLASTed to two databases: the *E. coli* K-12 genome (taxid 511145) and pSCL565, to capture spacers derived from the Cas1-Cas2 expression plasmid. To identify the source of acquired spacers during CRISPR-Cas primed adaptation amidst phage infection experiments, unique spacers extracted in steps described above were BLASTed to four databases: the *E. coli* K-12 genome (taxid 511145); the bacteriophage lambda genome (taxid 2681611); and pSCL787 or pSCL788, to capture spacers derived from the defence plasmids. In both cases, BLAST searches were performed with high stringency (≥90% identity, i.e., 30/33bp match between spacer and reference query) to obtain unique matches to the reference maps. We then parsed the BLAST results and filtered the genome-matching spacers for LacI, Cas1 and Cas2, as we assumed that spacers from these sources were most likely plasmid-derived.

#### Mapping spacers to reference genomes

Using the spacer genomic (lambda or *E. coli* K-12) or plasmidic (pSCL565, pSCL787 and pSCL788) location, target locus and counts, we generated coverage maps of the different genomes and plasmids where the spacers could have been sourced from, as well as spacer counts per location (i.e., counts of how many of the new spacers were *E. coli*, lambda, or plasmid derived). Briefly, for each BLAST record, we first checked whether the BLAST record mapped to any of our reference genomes, and if so, added counts (from **b.**) to spacer origin and occupancy counters. The occupancy array is generated analogously to those used to estimate sgRNA coverage (see **2.** above), and is genome-size aware (i.e., accounts for start-end junctions).

#### Spacer neighbourhood analysis

We also used the spacer →← genome information to look into the 15bp up and downstream of the genomic origin of the new spacer, in the hopes of capturing information regarding the PAM (canonically, AAG for this CRISPR adaptation system) and any other discernible motifs. This was done by mapping the spacer back to its reference genome, using the BLAST results, and extracting 15 bases upstream and downstream of the spacer. These sequences were compiled and Logomaker^85^ was used to generate sequence logos for the up and downstream region. This yielded **Fig 2d**.

#### Spacer origin distribution

Next, we used the spacer origin counters to obtain information about the breakdown of spacers by their origin (*E. coli*, lambda, or plasmids). To do so, we first normalised the spacer count per location to the number of arrays sequenced. In parallel, we also normalised the spacer count per location to the total number of new spacers identified, converting this metric to the percent of spacers mapping to each location. This allowed us to then plot the new spacer count with respect to spacer origin and strain of interest (**Fig 2b** and **Fig 5b**), in addition to the percent of new spacers belonging to each spacer origin and strain of interest (**Fig 2c**).

#### Coverage plots

Lastly, we generated genome coverage plots for *E. coli*, lambda, and the plasmids, as described in **2**. above. For the *E. coli* and lambda genomes, we generated binned coverage plots by calculating the coverage as a sliding mean, or binned coverage. The spacer coverage for plasmids was generated without binning spacer occupancy. This analysis yielded **Fig 2e-h**, **Fig 5c**, **Extended Data figure 1** and **Extended Data figure 5**.

The Jupyter Notebook for this analysis can be found here: https://github.com/Shipman-Lab/CRISPRi_host_factor_screen/blob/main/map_new_spacers_clean.ipynb

### Biological replicates

Biological replicates were taken from distinct samples, not the same sample measured repeatedly.

## Data Availability

All data supporting the findings of this study are available within the article and its supplementary information. Data used to generate all figures and perform statistical analysis, alongside a Jupyter Notebook to recreate our figures is available on GitHub here: https://github.com/Shipman-Lab/CRISPRi_host_factor_screen/blob/main/plot_run_stats_clean.ipynb. All sequencing data associated with this study is available on NCBI SRA (PRJNA1109382).

## Code Availability

All code used to process or analyse data from this study is available on GitHub here: https://github.com/Shipman-Lab/CRISPRi_host_factor_screen.

## Acknowledgements

Work was supported by funding from the National Science Foundation (MCB 2137692), the Pew Biomedical Scholars Program and the Chan Zuckerberg Biohub – San Francisco. S.C.L. was supported by a Berkeley Fellowship for Graduate Study. We thank Kate Crawford and Alex González Delgado for experimental guidance, assistance and comments on the manuscript.

## Author Contributions

S.C.L. conceived the study, and with S.L.S., outlined the scope of the project and designed experiments. All experiments were performed and analysed by S.C.L., with technical assistance from Y.L. and K.A.Z for Fig 1e (Y.L.) and Fig. 4f (K.A.Z.). S.C.L. and S.L.S. wrote the manuscript, with input from all authors.

## Extended Data Figures

**Extended Data Figure 1.**
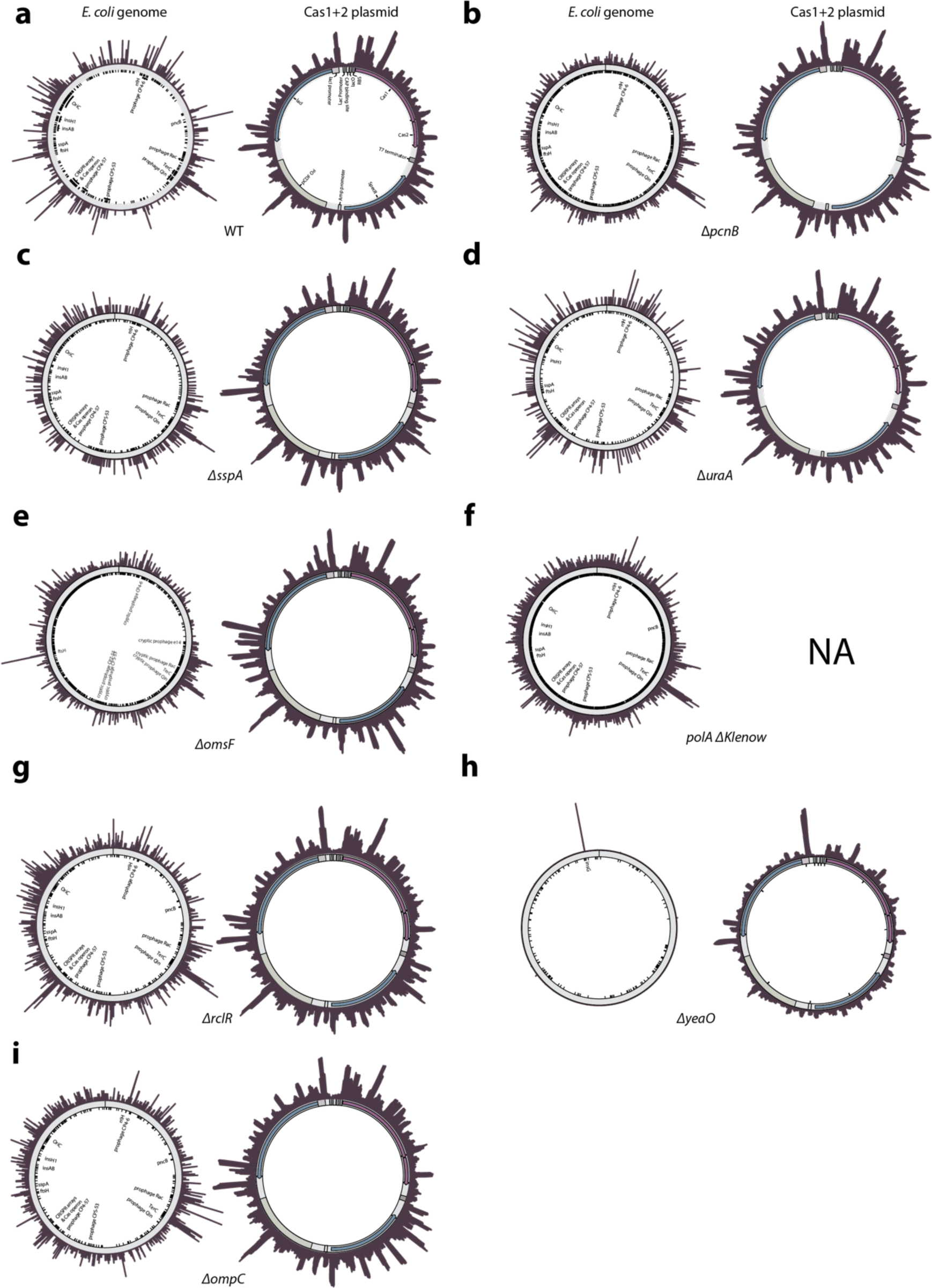
Binned coverage plot of newly acquired spacer across the *E. coli* genome (left) and pSCL565 plasmid (right) for strains selected for individual validation. **a-i**: wild-type, Δ*pcnB*, Δ*sspA*, Δ*uraA*, Δ*omsF*, *polA* ΔKlenow, Δ*rclR*, Δ*yeaO* and Δ*ompC*. Wild-type is *E. coli* BW25113, parental strain to the Keio collection; all other strains besides *polA* ΔKlenow are from the Keio collection. *polA* ΔKlenow was constructed as described previously^47^.

**Extended Data Figure 2.**
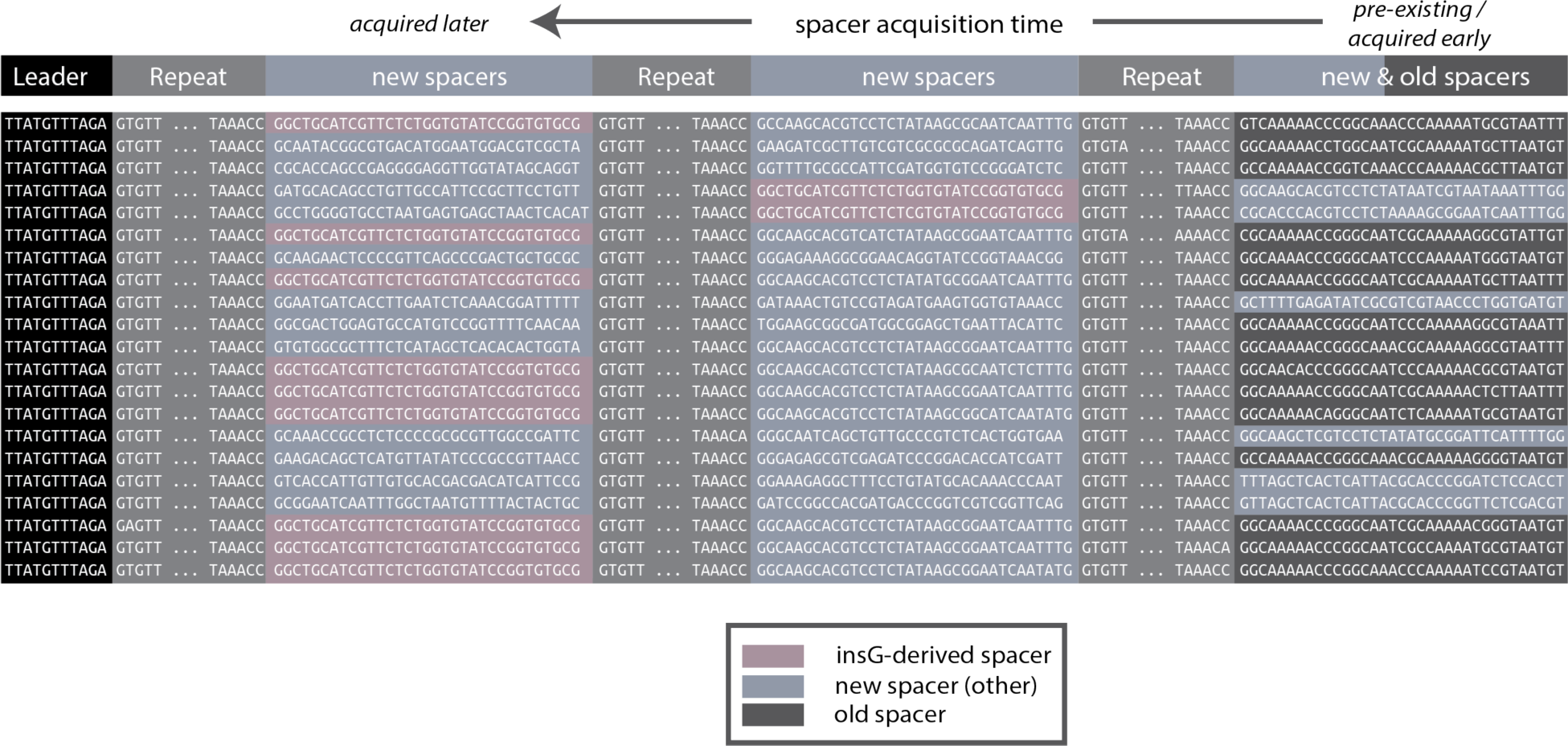
Alignment of sequencing reads corresponding to doubly and triply expanded CRISPR array from the Δ*yeaO* mutant strain, highlighting *insG*-derived spacers, new spacers and old or pre-existing spacers. with spacers acquired at a later stage are closer to the Leader sequence than spacers acquired earlier, or than pre-existing _spacers_49,51,52.

**Extended Data Figure 3.**
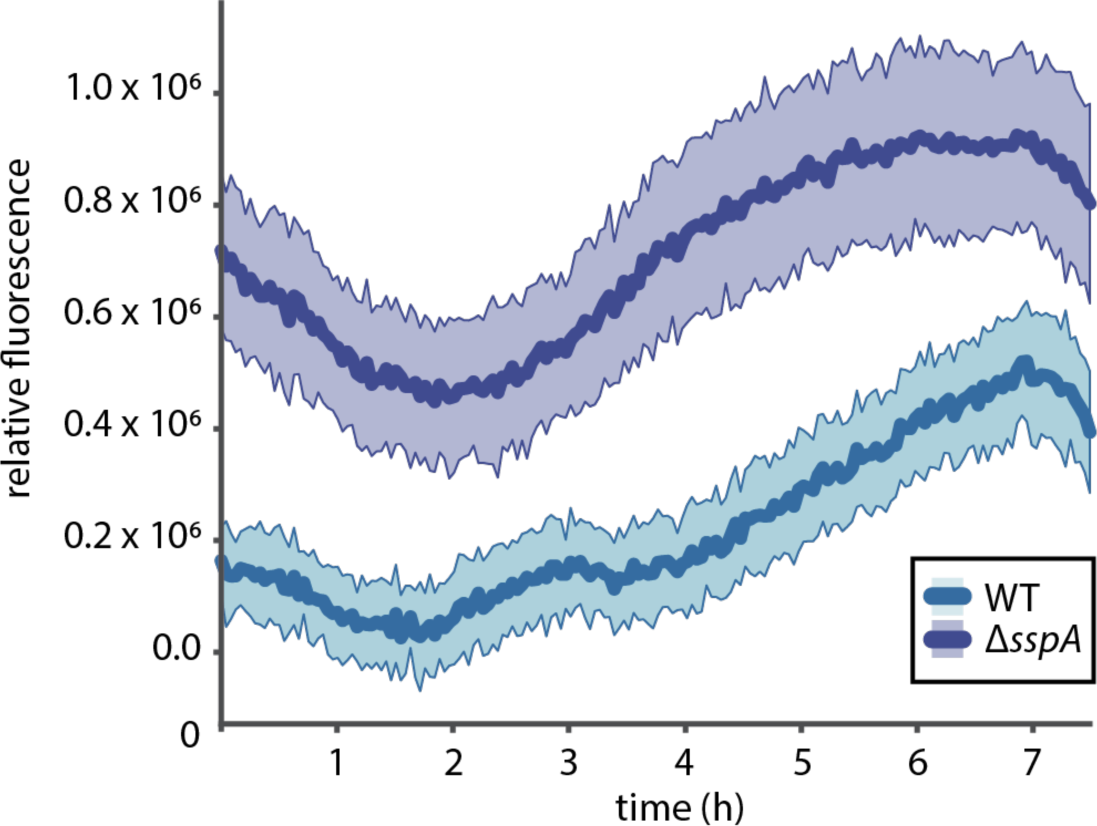
Fluorescence-based monitoring of the Lac promoter activity, used to express the Cas1-Cas2 integrases on pSCL565, in wild-type and Δ*sspA* cells, over the course of 7h of liquid culture. Hue around solid line (mean) represents the standard deviation across 3 biological replicates.

**Extended Data Figure 4.**
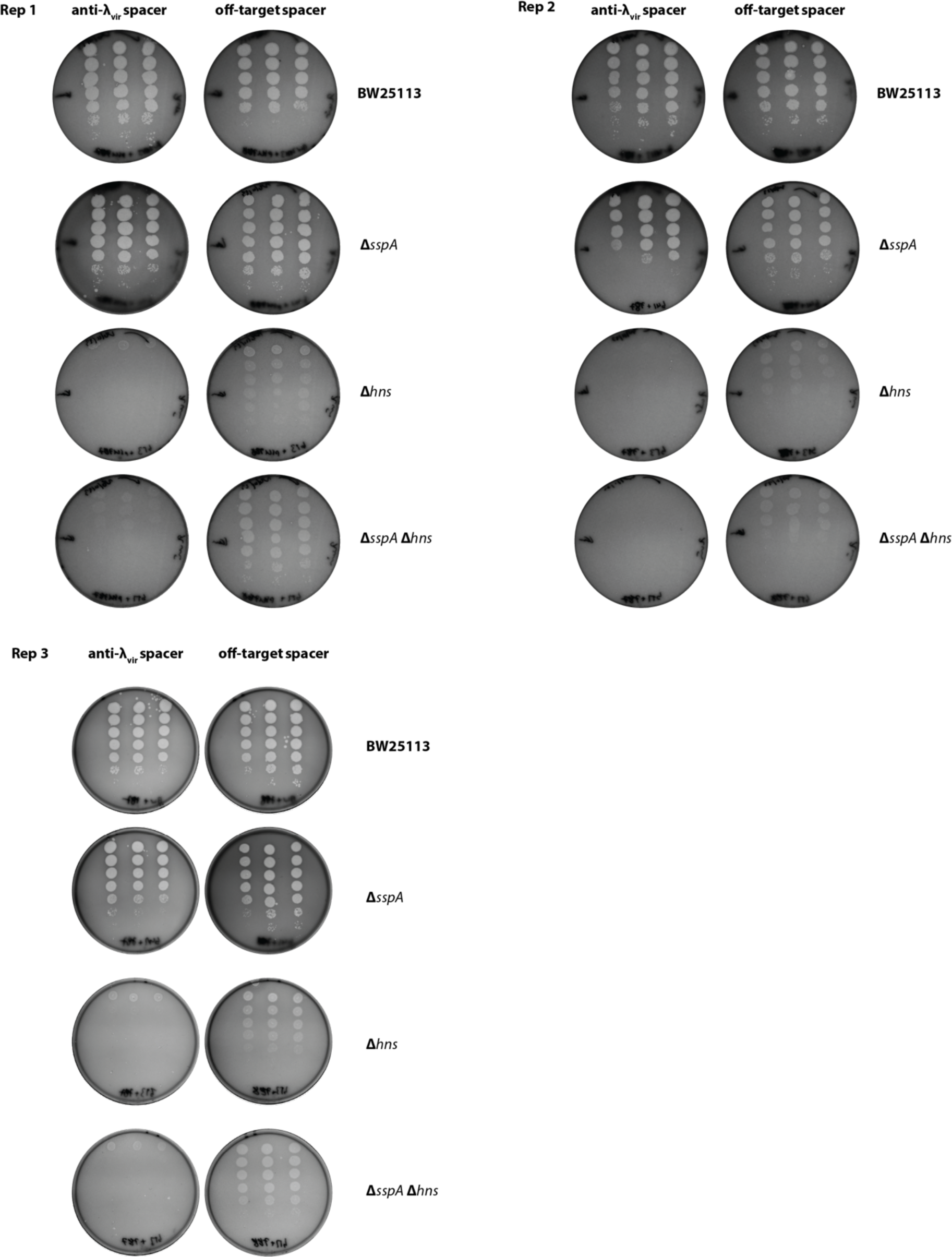
Full plates and 3 biological replicates of plaque assays of αvir on WT, Δ*sspA*::*FRT,* Δ*hns*::*FRT* and Δ*sspA*::*FRT* Δ*hns*::*FRT* strains, pre-immunised with either T or NT defence plasmids, corresponding to Fig 4d. Strains were infected with αvir and grown on plates at 30°C for 16h.

**Extended Data Figure 5.**
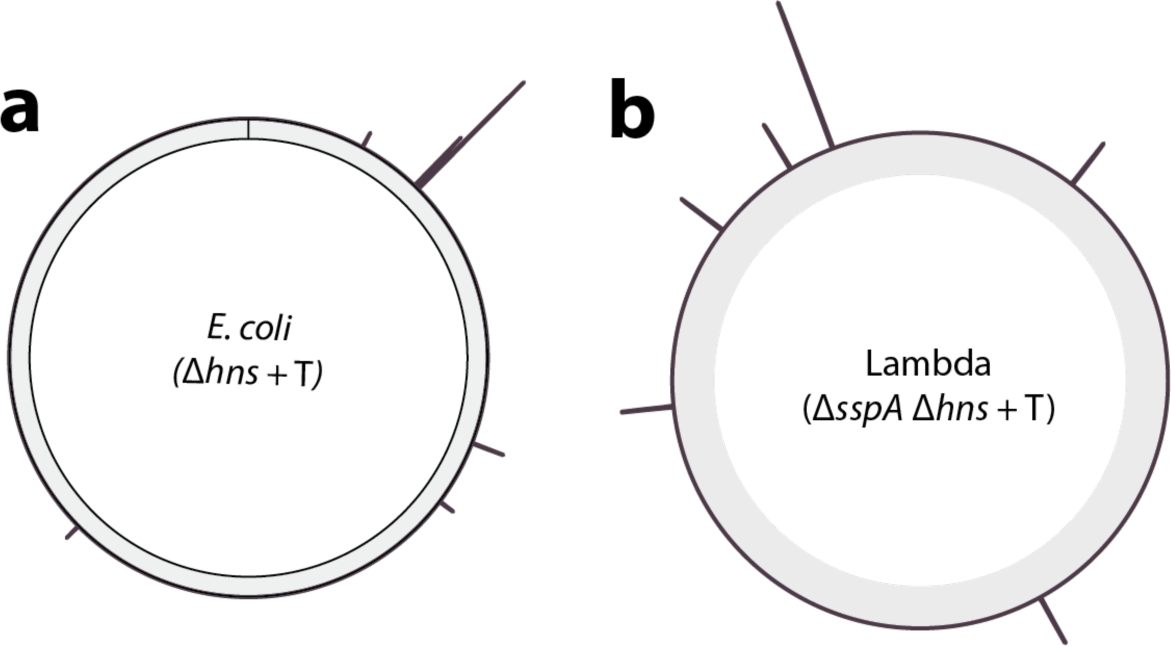
Distribution of newly acquired spacers in Δ*hns* +T and Δ*sspA* Δ*hns* +T strains upon lambda infection. **a**. Binned coverage plot of Δ*hns* + T newly acquired spacers across the *E. coli* genome (outer, purple). **b**. Binned coverage plot of Δ*sspA* Δ*hns* + T newly acquired spacers across the lambda genome (outer, purple).

